# Prosaposin maintains adult neural stem cells in a state associated with deep quiescence

**DOI:** 10.1101/2023.08.08.552453

**Authors:** Miriam Labusch, Melina Thetiot, Emmanuel Than-Trong, David Morizet, Marion Coolen, Hugo Varet, Rachel Legendre, Sara Ortica, Laure Mancini, Laure Bally-Cuif

## Abstract

In most vertebrates, adult neural stem cells (NSCs) continuously give rise to neurons in discrete brain regions. A critical process for maintaining NSC pools over long periods of time in the adult brain is NSC quiescence, a reversible and tightly regulated state of cell cycle arrest. Recently, lysosomes were identified to regulate the NSC quiescence-proliferation balance. However, it remains controversial whether lysosomal activity promotes NSC proliferation or quiescence, and a finer influence of lysosomal activity on NSC quiescence duration or depth remains unexplored. Using RNA-sequencing and pharmacological manipulations, we show that lysosomes are necessary for NSC quiescence maintenance. Additionally, we reveal that expression of *psap*, encoding the lysosomal regulator Prosaposin, is enriched in quiescent NSCs (qNSCs) that reside upstream in the NSC lineage and display a deep/long quiescence phase in the adult zebrafish telencephalon. We show that shRNA-mediated *psap* knock-down increases the proportion of activated NSCs (aNSCs) as well as NSCs that reside in shallower quiescence states (signed by *ascl1a* and *deltaA* expression). Collectively, our results identify the lysosomal protein Psap as a (direct or indirect) quiescence regulator and unfold the interplay between lysosomal function and NSC quiescence heterogeneities.

## 1. Introduction

Adult neural stem cells (NSCs) produce neurons within discrete regions of most vertebrate brains, including the mouse sub-ependymal zone of the lateral ventricle (SEZ), the sub-granular zone of the dentate gyrus of the hippocampus (SGZ), and the zebrafish pallium [1,2]. NSC maintenance is permitted, in part, by their residing in quiescence, a reversible and actively maintained state of cell cycle arrest corresponding, in that case, to the G0 phase of the cell cycle [3]. A high frequency of NSC cell cycle entry (referred to as NSC activation) is linked with NSC exhaustion [4–7]. Molecular pathways such as Notch2/3 signaling promote NSC quiescence in the different neurogenic domains, but single cell (sc) RNA-Seq, *in vivo* functional assays –e.g., the speed of NSC reactivation upon Notch signaling blockade and intravital imaging-, further show that quiescence is heterogeneous: NSCs can be found in shallower or deeper, or shorter or longer, quiescence [8–14]. Notably, a pre-activated state was identified by the expression of *Ascl1,* encoding a bHLH transcription factor important for the activation of previously quiescent NSCs [15–18]. Our recent results in the adult zebrafish pallium also highlight that the expression of *deltaA* (*deltaA^pos^*) correlates with NSCs that on average display shorter quiescence durations than *deltaA*-negative (*deltaA^neg^*) NSCs and are further advanced along the neurogenesis lineage [11]. Aside from these few examples, the cellular and molecular mechanisms underlying the control of quiescence depth/length remain incompletely understood.

Recently, attention has been given to differences in sub-cellular organelles between activated (cycling) NSCs (aNSCs) and quiescent NSCs (qNSCs). Among these, lysosomes emerged as novel regulators of NSC quiescence. Work in mice showed that qNSCs are enriched in lysosomes compared to aNSCs in the SGZ and SEZ [19,20]. However, these studies show discrepant results upon lysosomal perturbation. Impairment of lysosome activity through pharmacological inhibition of the vacuolar H(+)-ATPase with Bafilomycin A (BafA) leads to decreased activation of qNSCs in response to growth factors in the SEZ due to accumulations of protein aggregates [20]. In contrast, in the SGZ, BafA treatment leads to the accumulation of activated EGFR, Notch1, and the Notch1 intracellular domain NICD, increasing NSC activation [19]. Whether these differences are due to the different neurogenic niches or to different experimental designs remains undeciphered. Furthermore, next to their role in proteostasis maintenance, lysosomes act as important players in nutrient– and pathogen sensing, as well as membrane repair [21], and it remains to be studied whether the effect of lysosome perturbation on quiescence also involves these pathways. Finally, it is unknown whether lysosomal activity relates to quiescence heterogeneities between NSCs.

To address these issues, we focused on NSCs of the zebrafish adult pallium, which are easily accessible for transient functional manipulations *in vivo*, and for which we generated a comprehensive scRNAseq database and *in situ* readouts identifying qNSC heterogeneities correlating with differences in quiescence depth or length [11,13]. We report here that lysosomal activity is necessary for NSC quiescence in this system, and we identify the lysosomal protein Prosaposin (Psap) as a critical player in this activity. First, *psap* expression is highest in *deltaA^neg^* qNSCs, which display deepest/longest quiescence. Second, *psap* knockdown promotes qNSCs transition through the *deltaA^pos^* state of shallower/shorter quiescence, and NSC activation. These results reinforce the view that lysosomal activity promotes NSC quiescence and further highlight Psap as a likely player in this process.

## 2. Materials and Methods

### 2.1. Animals

Wild-type (AB), Tg(*gfap:dTomato*) [22], Tg(*gfap:egfp*) [23], Tg(4.5*deltaA:gfp*) (later referred to as Tg(*deltaA:gfp*) [24], Tg(*her4.3:drfp*) (later referred to as Tg(*her4:gfp*) [25], Tg(*mcm5:gfp*) [26], Tg(*her4:Venus-PEST*) [27] and Tg(*mcm5:nls-rfp*) [this study] zebrafish were maintained using standard protocols and in accordance with Institute Guidelines for Animal Welfare. Three-to four-month-old adult fish were used for analysis unless otherwise stated. Zebrafish were kept in 3.5-liter tanks at a maximal density of five per liter, in 28.5 C and pH 7.4 water.

### 2.2. Generation of the Tg(mcm5:nls-rfp)^ip^ transgenic line

To generate the *Tg(mcm5:nlsRFP)*^ip^ line, the multisite Gateway technology (Invitrogen^TM^) was employed with Tol2kit plasmids [28]. The *mcm5* promoter was first cloned from an in-house plasmid [29] using the following primers (forward: *5-GATCCTCGAGCATTCACCAGGGGTTCACAG-3’*; reverse: *5’-GATCGGATCCCTTGCAAATATTCAGGAATGTACAC-3’*) and the Invitrogen^TM^ Platinum Taq polymerase, then digested with XhoI and BamHI and ligated into the digested *p5E-MCS* backbone (Tol2kit #228) to obtain the *p5E-mcm5* plasmid. A fragment containing the first exon, the first intron and part of the second exon of the *mcm5* gene, followed in frame by the *P2A* [30] and *nlsRFP* sequences was synthetized by IDT, digested with XhoI and HindIII and ligated into *pME-MCS* to obtain the *pME-nlsRFP* plasmid. Finally, *p5E-mcm5*, *pME-nlsRFP* and *p3E-polyA* (Tol2kit #302) were recombined in the *pDestTol2CG2* (Tol2kit #395) vector using the Invitrogen^TM^ Gateway LR ClonaseII Plus enzyme to obtain the final plasmid. This plasmid was injected in 1-celled *casper* [31] or *AB* embryos together with 40ng/µl of *transposase* capped RNA. F0 adults crossed with *casper* fish were screened for transmission of fluorescence (green heart and red cells) in F1 embryos to generate a stable line.

### 2.3. Ventricular Microinjections and Electroporations

Micro-injections into the adult pallial ventricle were performed on anesthetized fish as described [32]. Morpholinos (MOs) (Gene-tools, Philomath, OR, USA) were injected at a concentration of 0,5 mM in PBS with 250 ng/µl of plasmid DNA. For plasmid electroporations, plasmid DNA was diluted to a final concentration of 500 ng/µl in PBS and injected into the ventricle. Fish were then administered four electric pulses (50 V, 50 ms width, 1,000 ms space) and were sacrificed after 3, 5, and 14 days.

### 2.4. RNAScope whole-mount in situ hybridizations

RNAScope is a commercial kit (© ACD, Multiplex kit). Whole brains were fixed overnight in 4% paraformaldehyde (PFA) in PBS and kept in 100% methanol at −20°C. Following rehydration, a pigment removal solution was applied for 10 min under light to bleach the brains. Following washes, the brains were kept in the Probe Diluent solution at 40°C and then incubated with the probe (1:50) overnight at 40°C. On the second day, the probe was washed off at room temperature (RT), followed by 30 min Amp1 incubation at 40°C, then washes followed by 30 min Amp2 incubation at 40°C, washes, 30 min Amp3 incubation at 40°C, and washes at RT. The brains were incubated at 40°C for 15 min with HRP, then washed at least 6 times. Afterward, the brains were incubated for 10 min at 40°C in TSA solution and finally incubated with Opal 650 (1:150 in TSA) for 30 min at 40°C. The solution was washed off; then the brains were incubated for 15 min in an HRP blocker at 40°C.

We quantified expression of *ascl1a* in individual NSCs by manually counting, using the Imaris software (Bitplane) in 3D, the fluorescent dots found in close proximity (< 1 μm) to the cell nuclei, labeled with H2amCherry. We considered that a cell was *ascl1a*^pos^ when at least 2 dots were visible. To quantify expression of *psap* (for Figure 2f), we did Z MaxProjection in ImageJ and then defined regions of interest (ROI) based on the ZO1 signal. We extracted the intensity sum in each cell. The highest value was set to 1 and the lowest to 0 to normalize (to compare different brains). We counted at least 100 cells per brain.

### 2.5. RNAi

To isolate the *psap* 3’ UTR, we performed 3’ RACE [33]. To design shRNAs, the miR designer by Thermofisher was used, and the resulting shRNAs were ordered as dsDNA blocks containing surrounding regions with restriction sites matched to the final plasmid, R73. The dsDNA blocks were sub-cloned in TOPO TA (Strataclone), then digested with XhoI and BamHI and added to a p3 entry plasmid –R73-, which can be used for cloning with the Gateway system and contained the sequence of an intron. R73 was cut with BglII and XhoI. This ensures that previously added shRNAs are not cut out of R73 but that multiple shRNAs can be added step by step through each round of cloning, in which shRNAs are chained. For *psap*, 9 shRNAs were chained. Our control construct contained 3 shRNAs. The following Geneblocks were ordered:

Psap01

atcgggatcctggaggcttgctgaaggctgtaTGCTGAGAACTCAAAGTTTGGACCCAGTTTTGGCCACTGACTGACTGGGTCCACTTTGAGTTCTcaggacacaaggcctgttactagcactcacatggaacaaatggcccagatctggccgcactcga gatcg

Psap02

atcgggatcctggaggcttgctgaaggctgtaTGCTGAGAACTCAAAGTTTGGACCCAGTTTTGGCCACTGACTGACTGGGTCCACTTTGAGTTCTcaggacacaaggcctgttactagcactcacatggaacaaatggcccagatctggccgcactcga gatcg

Psap03

atcgggatcctggaggcttgctgaaggctgtaTGCTGCACAGCTTGAGACTAAGTTGCGTTTTGGCCACTGACTGACGCAACTTACTCAAGCTGTGcaggacacaaggcctgttactagcactcacatggaacaaatggcccagatctggccgcactcg agatcg

Psap04

atcgggatcctggaggcttgctgaaggctgtaTGCTGTGTTCTGGAAGTATTCTGCTCGTTTTGGCCACTGACTGACGAGCAGAACTTCCAGAACAcaggacacaaggcctgttactagcactcacatggaacaaatggcccagatctggccgcactcg agatcg

Psap05

atcgggatcctggaggcttgctgaaggctgtaTGCTGTCTACAGCAATTCTCCTTTAGGTTTTGGCCACTGACTGACCTAAAGGAATTGCTGTAGAcaggacacaaggcctgttactagcactcacatggaacaaatggcccagatctggccgcactcga gatcg

Psap06

atcgggatcctggaggcttgctgaaggctgtaTGCTGTAAATTAGCGGTGACCATTCAGTTTTGGCCACTGACTGACTGAATGGTCCGCTAATTTAcaggacacaaggcctgttactagcactcacatggaacaaatggcccagatctggccgcactcga gatcg

Psap07

atcgggatcctggaggcttgctgaaggctgtaTGCTGTAAAGAGGGAGTATTCTGCTCGTTTTGGCCACTGACTGACGAGCAGAACTCCCTCTTTAcaggacacaaggcctgttactagcactcacatggaacaaatggcccagatctggccgcactcga gatcg

Psap08

atcgggatcctggaggcttgctgaaggctgtaTGCTGAGAAATGTTTACATCACTGGCGTTTTGGCCACTGACTGACGCCAGTGATAAACATTTCTcaggacacaaggcctgttactagcactcacatggaacaaatggcccagatctggccgcactcga gatcg

Psap09

atcgggatcctggaggcttgctgaaggctgtaTGCTGTAAAGAAGCACAATGACAGAAGTTTTGGCCACTGACTGACTTCTGTCAGTGCTTCTTTAcaggacacaaggcctgttactagcactcacatggaacaaatggcccagatctggccgcactcga gatcg

Control1

ggatcctggaggcttgctgaaggctgtaTGCTGTGATTATGATCTAGAGTCGCGGTTTTGGCCACTGACTGACCGCGACTCGATCATAATCAcaggacacaaggcctgttactagcactcacatggaacaaatggcccagatctggccgcactcgag

Control2

ggatcctggaggcttgctgaaggctgtaTGCTGTTATGATCTAGAGTCGCGGCCGTTTTGGCCACTGACTGACGGCCGCGACTAGATCATAAcaggacacaaggcctgttactagcactcacatggaacaaatggcccagatctggccgcactcgag

Control3

ggatcctggaggcttgctgaaggctgtaTGCTGTTATGATCTAGAGTCGCGGCCGTTTTGGCCACTGACTGACGGCCGCGACTAGATCATAAcaggacacaaggcctgttactagcactcacatggaacaaatggcccagatctggccgcactcgag

To validate knock-downs, we used RNAScope and quantified the RNA signal in transfected cells, as follows. We did Z MaxProjection in ImageJ and then defined regions of interest thanks to the ZO1 signal. Per transfected cell, we selected 4 neighboring cells to be able to normalize and measure the intensity sum in electroporated vs. non-electroporated cells of the RNAScope signal. We then divided the average signal in electroporated cells by the average signal in non-electroporated cells to measure expression changes.

### 2.6. Organotypic slices generation and BafA treatment

3-mpf zebrafish brains were dissected in DMEM/F-12 with GlutaMax (11514436, Thermo Fisher Scientific), complemented with penicillin-streptomycin (100 IU/ml, 15140122, Thermo Fisher Scientific). They were cut into 200 µm coronal slices using a McIlwain tissue chopper, and the slices were placed on Millicell membranes (3–4 slices per membrane, 0,4 µm Millicell, Merck Millipore). Slices were cultured in DMEM/F-12 with GlutaMax, B-27 supplement (2%, 17504044, Thermo Fisher Scientific), N2 supplement (1%, 17502048, Thermo Fisher Scientific), Insulin solution human (4 µg/ml, I9278, Sigma), D-Glucose (0,3%; G8769, Sigma), amphotericin B (0,10 µg/ml, 15290026, Thermo Fisher Scientific) and penicillin-streptomycin (100 IU/ml, 15140122, Thermo Fisher Scientific), and supplemented with EGF (20 ng/ml, E4643, Sigma) and FGF2 (20 ng/ml, 100-18b, Tebu-bio). Cultures were maintained at 28.5°C under 5% CO2 perfusion. To chemically block the v-ATPase, slices were incubated for 1 day in vitro (DIV) in either BafA (24h, 20 nM, SML1661, Sigma-Aldrich) or DMSO (24h, 0.02%, D8418, Sigma-Aldrich) added to the fresh culture medium. To evaluate the impact of lysosomal deacidification, slices were fixed 24h post-treatment in 4 % PFA for 30 min at RT and washed with PBT, 3 times 10 min. Slices were then detached from the membranes for immunohistochemistry.

### 2.7. Quantification of lysosomes in NSCs

Dissected pallia (experiment: 10 pallia; controls: 5 pallia each) were recovered and pooled in PBS and then transferred into pre-warmed FACSmax solution (AMS.T200100, Amsbio), in which the pallia were incubated for 4 min at 28.5°C, then placed on a 40 µm nylon cell strainer (BD Falcon^TM^) and gently passed through it with a 1 ml syringe plunger. The dissociated cell solution was recovered into a Petri dish and transferred to a 15 ml Falcon tube. The strainer and Petri dish were washed with FACSmax solution to increase the number of recovered cells, which were also added to the Falcon and mixed by gentle pipetting. Dissociated tissues from adult brains were centrifuged at 500 g for 5 min. After discarding the supernatant, cells were incubated in a lysotracker solution (lysotracker, L12492, Invitrogen, 1:10 000 in PBS) for 4 min at 28.5°C and then centrifuged at 500 g for 3 min. The supernatant was discarded, and the cells were washed with PBS, then centrifuged for 3 min at 500 g, and finally recovered in PBS + 1 mM EDTA + 1 µg/ml DAPI (4’,6-diamidino-2-phenylindole). DAPI was used to remove dying cells from the analysis later on. Flow cytometry was performed using the Becton Dickinson FACSymphony A5. As controls, every fluorophore was measured by itself (AB with DAPI, Tg(*gfap:egfp*), Tg(*mcm5:nls-RFP*), AB with lysotracker) through flow cytometry. A minimum of 8 x 10^5^ events was recorded.

### 2.8. Transcriptomic analysis of telencephalic NSCs

#### Cell dissociation and Fluorescence-activated cell sorting (FACS)

*Tg(her4:dRFP)*/+;*Tg(mcm5:GFP)*/+ double heterozygote transgenic fish were euthanized in ice-cold water and their brain dissected in Ringer’s solution. Their telencephala were then isolated and collected into 2 ml tubes filled with 300 µl of cell dissociation solution (FACSmax^TM^) and kept on ice. The telencephala were then transferred to a 40 µm nylon cell strainer (BD Falcon^TM^) previously moistened with FACSmax^TM^ and the tissues were gently passed through the strainer (Falcon or BD Plastipak 300013) with the plunger of a 1 ml syringe. The dissociated cell solution was recovered into a 35 × 10 mm tissue culture dish (Falcon) placed on ice, pipetted back to the 2 ml tube, and mixed by gentle pipetting. This cell suspension was centrifuged at 1000 g for 4 min at 4°C. After discarding the supernatant, the cell pellet was resuspended with 600 µl of DMEM/F12 supplemented with 0.25X B27, 1X N2, glucose, and 20 µg/ml insulin, and containing 1 µg/ml DAPI ((4’,6-diamidino-2-phenylindole) and placed on ice until sorting. DAPI was used to label dying cells to remove them upon sorting. Ultimately, each sample comprised between 27 and 40 pooled telencephala.

Cells were directly sorted into 1.5 ml tubes containing 100 µl of extraction buffer (XB, Arcturus PicoPure RNA isolation kit) with a FACSAria III SORP (BD Biosciences) flow cytometer equipped with a 90 µm-diameter nozzle. PBS was used as a sheath liquid. To avoid an excessive dilution of the extraction buffer, a maximum of 20,000 events (i.e., cells) were sorted in each tube, often resulting in the collection of several tubes of each cell type.

Dissociated cells from the telencephala of AB, *Tg(her4:dRFP)*/+, and *Tg(mcm5:eGFP)*/+ fish were used in pilot experiments to gate eGFP and dRFP fluorescence. After excluding cell debris on FSC-A/SSC-A scatter plots, single cell events (singlets) were selected on both FSC-A/FSC-W and SSC-A/SSC-W plots, and dead or dying cells were excluded based on their positivity for DAPI. Finally, qNSC (dRFP^pos^/GFP^neg^), aNSCs (dRFP^pos^/GFP^pos^), a proliferating cell population comprising some proliferating neural progenitors (dRFP^neg^/GFP^pos^) and the remaining cells (dRFP^neg^/GFP^neg^) were selected and directly sorted into the extraction buffer. In addition, we also carried out another sorting experiment in which we distinguished qNSCs based on their level of *her4.3* expression: qNSC (*her4^high^*) (dRFP^high^/GFP^neg^) and qNSC (*her4^low^*) (dRFP^low^/GFP^neg^).

Overall, 3 independent sorting experiments were carried out for the comparison of juvenile and adult NSCs, and 4 independent experiments were performed for the comparison of *her4^high^* qNSCs, *her4^low^* qNSCs, and aNSCs.

#### RNA isolation

Total RNA was extracted and purified with the Arcturus PicoPure Isolation kit, according to the manufacturer’s protocol (Life Technologies). When sorting a specific cell type resulted in the collection of several tubes (see above), the ensuing extracted RNA samples were pooled just after their precipitation in 70% ethanol by binding them consecutively on the same purification column. An on-column DNA removal step was systematically carried out with RNase-free DNase Set (Qiagen®), according to Arcturus PicoPure Isolation kit protocol. Purified total RNA was stored at –80°C until further processing. The integrity of RNA samples was assessed on an Agilent 2100 Bioanalyzer, using an Agilent RNA 6000 Pico Kit. RNA integrity numbers (RIN) obtained were comprised between 8.3 and 9.10.

#### Library construction and RNA-sequencing

Libraries were constructed with the SMARTer® Stranded Total RNA-Seq Kit – Pico Input Mammalian kit (Clontech Laboratories), according to the manufacturer’s instructions. Briefly, RNA was fragmented for 4 min (RINs>7) and cDNA was synthesized using a mixture of random primers attached to the reverse PCR1 primer. The forward PCR1 primer was included in a locked nucleic acid-template-switching oligonucleotide (LNA-TSO) able to base-pair with the few nucleotides added to the 3’ end of the cDNA by the SMARTScribe™ Reverse Transcriptase (RT)’s terminal transferase activity. The extended template created by the LNA-TSO enables the RT to continue replicating to the end of the

oligonucleotide and to incorporate the forward PCR1 primer at the 3’ end of the cDNA. Full-length Illumina adapters, including barcodes (i.e., indexes, 8 nucleotide-long), were added by PCR amplification (PCR1). Ribosomal cDNA (originating from rRNA 18S and 28S) were then depleted through digestion by ZapR after hybridization to R-Probes. Finally, these non-digested fragments were amplified by PCR (PCR2), using primers universal to all libraries. Libraries concentrations and size distributions were assessed by running samples on Agilent 2100 Bioanalyzer using the Agilent High Sensitivity DNA Kit (Agilent, Part Number 5067-4626). cDNA sample size modes were about 300 nucleotides (N.B.: library preparation adds 139 bp to the size of the original cDNA fragments). More precise and accurate concentration estimates were obtained with a fluorescent-based quantitation assay (QuBit fluorometer, “Quant-It” assays kit, Invitrogen). Concentrations were normalized to 2 nM, and samples were multiplexed, denatured with 0.1N NaOH, and diluted to a final concentration of 8 pM before being loaded on the flowcell. Libraries were sequenced on a HiSeq 2500 Illumina sequencing machine as single reads, with 65 cycles. Hiseq SR Cluster Kit V4 and Hiseq SBS Kit-HS kits were used for cBot cluster amplification and sequencing, respectively.

#### Mapping and quality control of the reads

Reads were mapped to zebrafish genome assembly GRCz10 using the STAR (Spliced Transcripts Alignment to a Reference) aligner, thus accounting for splice variants. The quality of the reads was assessed with FastQC and revealed overall good quality, no contamination, effective rRNA depletion, and good mapping despite a high multimapping rate. Reads with multiple alignments were kept in the analysis.

#### Differential expression analysis

The analysis was performed using the R software, and Bioconductor packages including DESeq2. Normalization and differential analysis were carried out according to the DESeq2 model and package (version 1.12.3). The replicate effect was taken into account in the statistical models. Overall, 26,801 and 30,512 features were detected in the her4^high^/her4^low^/aNSCs and in the juvenile/adult NSCs comparisons, respectively. The number of mapped reads varied between 1 and 16 million across samples and replicates and accounted for 62-96% of all features. Read counts were normalized in DESeq2 by dividing them with a scaling factor associated with the sample they belong to and computed with locfunc=“shorth”. To determine the shape of the mean-variance relationship, the dispersion of the data was estimated in DESeq2 using a Generalized Linear Model (GLM)-based method (fitType=“parametric”). To increase the detection power of differentially expressed features at the same experiment-wide type I error, normalized read counts were pre-filtered by applying a threshold based on their mean, irrespective of the biological condition. Adjusted p-values of discarded features were then set to NA. To account for multiple testing and to control for the false discovery rate, p-values were adjusted according to the Benjamini–Hochberg’s method, and the significance level for the false discovery rate (FDR) was set at 0.05.

#### Normalized expression values

To make expression values comparable between features, read counts were first normalized with DESeq2 (to make reads comparable between samples), then divided by the feature length (in nucleotides), and finally multiplied by 1000 to obtain expression levels by kilobase.

#### Pathway enrichment analysis

To identify the pathways enriched among the genes significantly upregulated or downregulated (FDR<0.05) between two given cell types, we used the functional annotation chart module of the Database for Annotation, Visualization and Integrated Discovery (DAVID). Pathway annotations were taken from the KEGG database. A threshold of 5 was applied on gene counts in such a way that only terms harboring at least 5 enriched genes were considered in the analysis. The cutoff for EASE score was set at 0.05. Data are displayed as the enrichment score [-log10(p-value)] of the KEGG pathways displaying an EASE<0.05. The number of genes (of the differentially expressed gene list assessed) associated with an enriched term is indicated above the bar corresponding to the enrichment score.

#### Geneset enrichment analysis (GSEA)

The datasets (i) “deeply” qNSCs (*her4^high^*) (dRFP^high^/GFP^neg^) vs. aNSC (dRFP^pos^/GFP^pos^) and (ii) adult qNSCs 3.5 mpf (dRFP^pos^/GFP^neg^) vs. juvenile qNSCs 1.5 mpf (dRFP^pos^/GFP^neg^) were analyzed. Geneset enrichment analysis was performed using GSEA software 4.1.0. Zebrafish ensemble gene IDs were converted to human gene symbols, by collapsing the dataset to gene symbols with a dedicated “chip” file. We used the gene sets associated with the KEGG canonical pathways available from the c2.cp.kegg.v7.5.1 database of the Broadinstitute. The background was obtained by permutating 1000 times the gene sets. Gene sets larger than 500 and smaller than 15 were excluded. The metric used for ranking gene was Signal2Noise and the enrichment statistic was normally weighted. An FDR < 0.25 was used as a cutoff. Enrichment score plot and heatmap of expression level of GSEA-enriched genes belonging to the leading-edge subset of a GSEA geneset were generated with the GSEA software 4.1.0.

### 2.9. Morpholinos

Morpholinos were ordered from Genetools. Sequences are as follows:

*sort1a:* 5’AATGTTTCGACCTACCGTGTGAGT3’

*sort1b:* 5’TTGGTGCAAGTGAACTTACGTTGT

Standard control: 5’CCTCTTACCTCAGTTACAATTTATA

### 2.10. Immunohistochemistry

Whole brains were fixed overnight in 4 % paraformaldehyde in PBS and kept in 100% methanol at −20°C. Following rehydration, an antigen-retrieval step, in which brains were incubated in Histo-VT One (Nacalai Tesque) for 60 min at 65°C, was performed for nuclear immunolabelling, such as Sox2 and PCNA. Alternatively, and specifically for BrdU stainings, brains were incubated in 2N HCl solution (Sigma-Aldrich, 258148) for 30 min at RT. Brains were kept in the blocking solution (5 % Normal Goat Serum, 0.1 % Triton-X, 0,1 % DMSO in PBS) (Sigma Life Science, 1002135493) for at least 1 hour before incubating in the primary AB containing-blocking solution overnight at 4°C. On the second day, brains were washed at least 6 times with PBT and then incubated overnight at 4°C in a blocking solution containing the secondary ABs. On the third day, brains were washed 3 times before imaging. Primary Antibodies were as follows:

Anti-GFP: polyclonal, chicken, IgY, 1:500, Aves Labs #GFP-1020

Anti-Sox2: monoclonal, mouse, IgG1, 1:250, Abcam #ab171380

Anti-Sox2: polyclonal, rabbit, IgG, 1:250, Abcam #ab97959

Anti-Pcna: polyclonal, rabbit, IgG, 1:500, GeneTex #GTX124496

Anti-dsRed: polyclonal, rabbit, IgG, 1:300, Clontech/Ozyme #632496

Anti-ZO1: monoclonal, mouse, IgG1, 1:200, Invitrogen #AB-16.0425

Secondary antibodies raised in goat coupled to AlexaFluor dyes (Invitrogen) were used (1:1000).

### 2.11. Imaging, Cell Counting, and Statistics

Images were taken using Zeiss LSM700 and LSM710 confocal microscopes using the 40x (oil) objectives. Images were processed using the ZEN (Zeiss) and Imaris (Bitplane) softwares. Cell counts on zebrafish brains were carried out on whole mount pallia. For electroporations, we only counted brains that had at least 30 successfully transfected NSCs. Imaris was used for all counts. Statistical analysis was carried out for all quantitative experiments using Prism and InVivoStat [34]. The normality of the residuals of the responses was assessed using normality probability plots. When data displayed an approximately Gaussian distribution, parametric tests were performed. We did not assume equal standard deviations, and therefore turned to t-tests with Welch’s corrections. Exceptions are the lysotracker experiments, where we normalized the data and therefore expected similar Standard deviations in biological replicates. For data that did not display a Gaussian distribution, we turned to nonparametric Mann-Whitney tests. Experimental data are expressed as median ± IQR.

## 3. Results

### 3.1. Lysosomes are necessary for NSC quiescence

Adult pallial NSCs in zebrafish and mice are radial glial cells expressing glial fibrillary acidic protein (Gfap) [35,36]. Transgenic reporter lines for *gfap* but also for the progenitor gene *her4.3* characterize the same population of progenitors in the adult zebrafish brain [37]. At any given time, only about 5-10 % of NSC are expressing the cell cycle marker Proliferating cell nuclear antigen (Pcna), and are therefore defined as activated [27]. By exclusion, other NSCs are considered quiescent. Because NSCs in different (sub)states are intermingled *in situ*, combined markers are needed to distinguish between activated NSCs (aNSCs: *gfap^pos^* or *her4^pos^*, Pcna^pos^), pre-activated qNSCs (*gfap^pos^* or *her4^pos^*, Pcna^neg^, *ascl1a*^pos^), and, among other qNSCs, those in shallow/shorter quiescence (*deltaA^pos^* qNSCs: *gfap ^pos^* or *her4^pos^*, Pcna^neg^, *deltaA^pos^*) and those in deep/longer quiescence (*deltaA^neg^* qNSCs: *gfap ^pos^* or *her4^pos^*, Pcna^neg^, *ascl1a*^neg^, *deltaA^neg^*) (Figure 1a).

**Figure 1.**
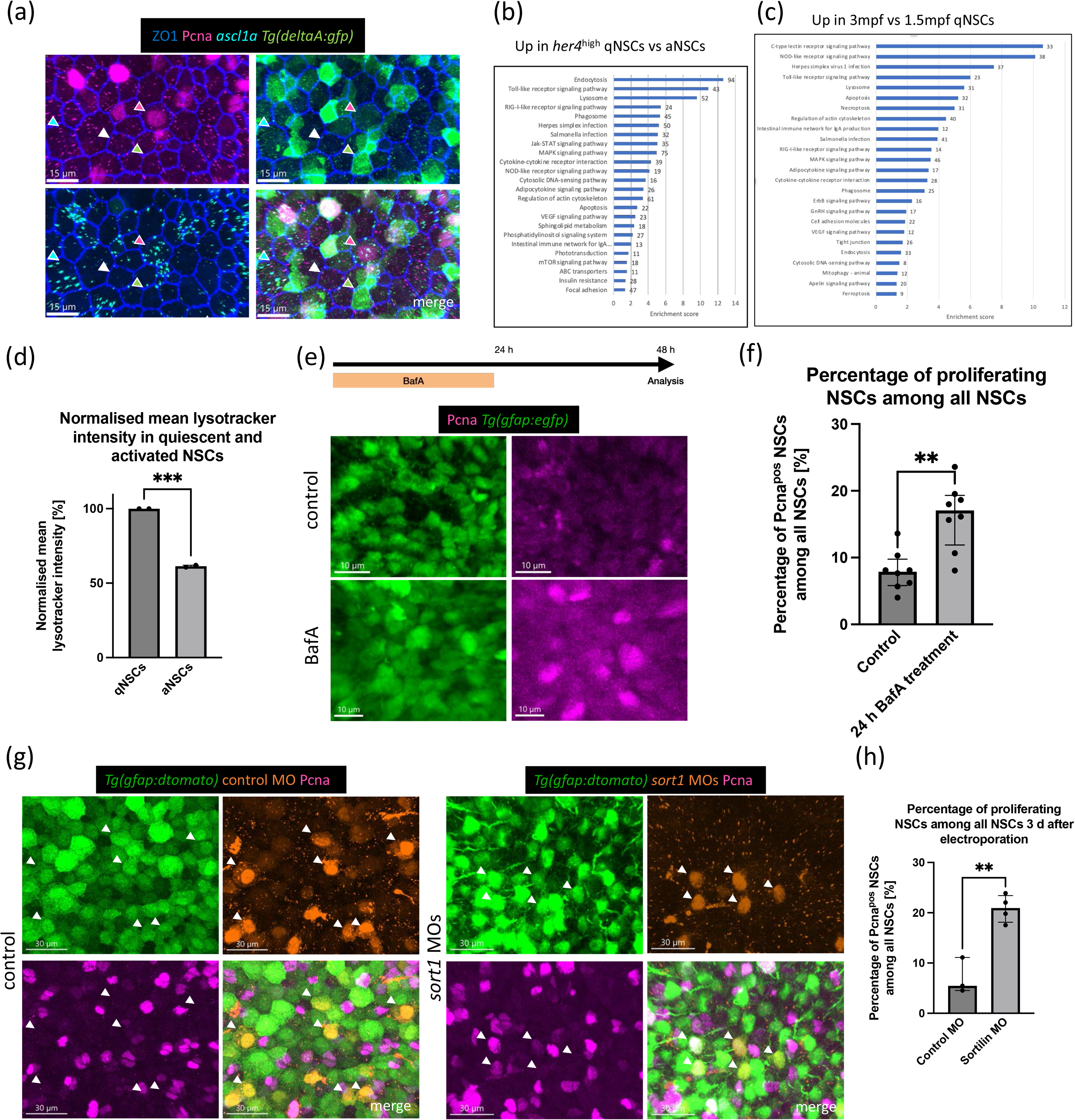
Lysosomes are critically involved in adult neural stem cell quiescence. **(a)** Whole mount immunostaining for ZO-1 (tight junctions, blue), Pcna (proliferation, magenta) and GFP (transcriptional reporter for *deltaA*, green) combined with RNAScope against *ascl1a* transcripts (pre-activation, cyan). Dorsal “apical” view of the pallial ventricular zone at 3 mpf. Green arrow points to *deltaA^pos^* qNSC, white arrow to *deltaA^neg^* qNSC, cyan to pre-activated NSC, and magenta to aNSC. Scale bar: 15 µm. **(b)** KEGG pathways enriched in qNSCs vs. aNSCs based on gene ontology analysis of the RNASeq data (RFP^high^,GFP^neg^ vs RFP^pos^,GFP^pos cells^ from 3mpf Tg(*gfap:egfp*);Tg(*mcm5:nls-rfp*) fish). On the x-axis, the enrichment score is represented. The number next to the bars gives information about the number of genes included in this pathway. The lysosomal pathway is highly enriched in qNSCs vs. aNSCs. **(c)** KEGG pathways enriched in qNSCs at 3.5 months vs. qNSCs at 1.5 months based on gene ontology analysis of the RNASeq data. On the x-axis, the enrichment score is represented. The number next to the bars gives information about the number of genes included in this pathway. The lysosomal pathway is enriched in older fish. **(d)** Normalized mean lysotracker intensity, represented in a barplot with SD. Each dot represents one independent experiment with 10 pooled zebrafish brains. aNSCs have a lower lysotracker intensity than qNSCs. Unpaired t-test, p = 0.0003. **(e)** Experimental design: Lysosomes were deacidified through BafA for 24 h (with 24 h recovery). View on organotypic slices after immuno staining for GFP (NSCs, green), and Pcna (proliferation, magenta) after control and BafA treatment. Scale bars: control 10 µm, BafA treated: 10 µm. **(f)** Proportion of proliferating NSCs among all NSCs. Each dot represents one animal (with 1-4 treated slices per animal). Bar at median with IQR. Mann-Whitney test, p = 0.0026. Proliferation is increased after BafA treatment compared to control treatment. **(g)** Whole mount immunostaining for Tg*(gfap:dTomato)* (NSCs, green), and Pcna (proliferation, magenta) in combination with control– and *sort1a/sort1b-*MO-electroporated cells (orange). Dorsal view of the pallial ventricular zone at 3 mpf. Arrows point to electroporated NSCs. Scale bar: 30 µm. **(h)** Percentage of activated NSCs among all electroporated NSCs 3 d after control– and *sort1*-MO electroporation. Each dot represents one animal. Line at median with IQR. Unpaired t-test with Welch’s correction, p = 0.0064. *sort1* MOs increase the percentage of aNSCs significantly.

To identify pathways regulating NSC quiescence, with a possible link to quiescence depth, we used double transgenic Tg(*her4:drfp*);Tg(*mcm5:gfp*) fish and two paradigms predicted to highlight deep vs shallower quiescence. First, in adult fish (3 month-post-fertilization –mpf-), we performed bulk RNA sequencing on FACS sorted RFP^high^,GFP^neg^ qNSCs and RFP^pos^,GFP^pos^ aNSCs. *her4.3* (Figure S1a, see [38]) is a target of Notch signaling, which promotes quiescence [27,39]. We identified 3048 genes upregulated in RFP^high^ qNSCs and 1890 genes upregulated in aNSCs (Tables S1 and S2). As expected for aNSCs, geneset enrichment analysis (GSEA) confirmed the enrichment for KEGG (Kyoto encyclopedia of genes and genomes) genesets associated with cell division, which include cell cycle (Figure S1b) and DNA replication (Figure S1c), as well as ribosomal activity (Figure S1d), as aNSCs exhibit higher protein synthesis than qNSCs [10]. Focusing on proteostasis, we observed enrichment of the proteasomal pathway in aNSCs (Figure S1e) and enrichment of the lysosomal pathway (Figure S1f) and the lysosome-related pathways autophagy (fig. S1Af) and endocytosis (Figure S1g) in RFP^high^ qNSCs. When operating gene ontology (GO) analysis, we identified the KEGG lysosomal pathway as highly enriched in RFP^high^ qNSCs (Figure 1b). Second, we also generated a dataset in which we compared the bulk transcriptome of qNSCs between 1.5 and 3.5 mpf fish, stages between which deeper quiescence is progressively instated (Tables S3 and S4). GO analysis identified the KEGG lysosomal pathway as significantly enriched at 3.5 mpf (Figure 1c), pointing to a pathway that becomes more prominent with time, as previously suggested between adult and aged mice [20].

Together, these data suggest that qNSCs may harbor more lysosomes than aNSCs, or increased lysosomal activity, or both. To address this, we performed flow cytometry analysis of freshly dissociated lysotracker-stained cells from adult pallia dissected from double transgenic Tg(*gfap:egfp*);Tg(*mcm5:nls-rfp*) fish, allowing us to dissociate between GFP^pos^; RFP^neg^ qNSCs and GFP^pos^; RFP^pos^ aNSCs (Figure S2). Lysotracker probes are fluorophores linked to a weak base that is only partially protonated at neutral pH and will emit fluorescence in acidic compartments, therefore in lysosomes [40]. We detected a decrease to 61.34 % (Median; IQR 60.70-61.99 %) in the normalized mean intensity of lysotracker fluorescence in aNSCs compared to qNSCs (Figure 1d), suggesting that qNSCs harbor more or enlarged lysosomes. We next wondered whether these lysosomes might be involved in quiescence maintenance. To impair lysosomal activity, we incubated organotypic slices of adult Tg(*gfap:egfp*) zebrafish pallia in 20 nM BafA for 24 h followed by 24 h of recovery (Figure 1e). Quantifying Pcna^pos^ GFP^pos^ aNSCs vs. Pcna^neg^ GFP^pos^ qNSCs revealed a significant percentual increase of aNSCs upon BafA treatment vs. control treatment (Median and IQR: control 7.89 %, 5.81-9.76 % vs. BafA treated 17.06 %, 11.90-19.32 %) (Figure 1f). Overall, we conclude that lysosomal content is enriched in the quiescent vs. activated NSC states and that lysosomal activity is critically involved in the maintenance of quiescence.

Next, to better understand whether quiescence control involves the proteostasis activity of lysosomes [21], we focused on Sortilin1, a sorting receptor of the vacuolar protein sorting 10 family that facilitates lysosome-mediated protein degradation. Sortilin1 is ubiquitously expressed in many tissues but most abundant in the central nervous system [41]. The zebrafish duplicates *sort1a* and *sort1b* were expressed together at a high level in qNSCs in our bulk dataset, and also detectable in our single cell NSC dataset of the adult zebrafish pallium [38] (Figure S3). To knockdown Sort1 function, we used previously validated splice-morpholino oligonucleotides (MOs) directed against *sort1a* and *sort1b* [42], which we electroporated *in vivo* into adult NSCs in Tg*(gfap:dTomato*) adult fish following intracerebral injection. We found that abrogating Sort1 function led to a significant increase in the proportion of aNSCs among cells electroporated with *sort1a*/*sort1b* MOs compared to control MOs (median and IQR: control 5.45 %, 4.55-11.11 % vs. *sort1a*/*sort1b* MOs 20.93 %, 18.11-23.42 %) (Figures 1g and 1h), reaching levels very similar to those obtained upon BafA treatment. Overall, we conclude that lysosomes are enriched in qNSCs that exhibit deep/long quiescence, and that their proteostasis activity promotes NSC quiescence in the adult brain *in vivo*.

### 3.2. psap is heterogeneously expressed in NSCs during their quiescence phase

To further challenge whether NSC quiescence involves proteolysis control by lysosomes, as proposed [19,20], and to unravel through which molecular players, we analyzed the expression of the genes that contributed to the enrichment of the lysosomal pathway in qNSCs in our RNAseq datasets. This led us to select *psap*, a gene encoding a glycoprotein involved in the activation of hydrolases in the lysosomes [43], and to date solely implicated in lysosomal proteostasis. In addition, cells deficient for Sort1 exhibit impaired trafficking of Psap to the lysosomes and enhanced release of Psap into the extracellular medium [44–47]. We found that *psap* was expressed at levels 3.5 times higher in qNSCs vs. aNSCs, in a manner very reproducible among replicates (Figure 2a). It is also enriched 2.3 times in 3mpf compared to 1.5mpf qNSCs (Table S3). RNAScope ISH experiments for *psap* further revealed heterogeneous expression levels among NSCs (Figure 2b). Finally, and importantly, *psap* transcript levels in individual qNSCs in our scRNAseq dataset were stronger in cells transcriptionally distant from *pcna-*expressing aNSCs, *ascl1a*-expressing pa-qNSCs and *deltaA*^pos^ NSCs (Figures 2c and S4a-c).

**Figure 2.**
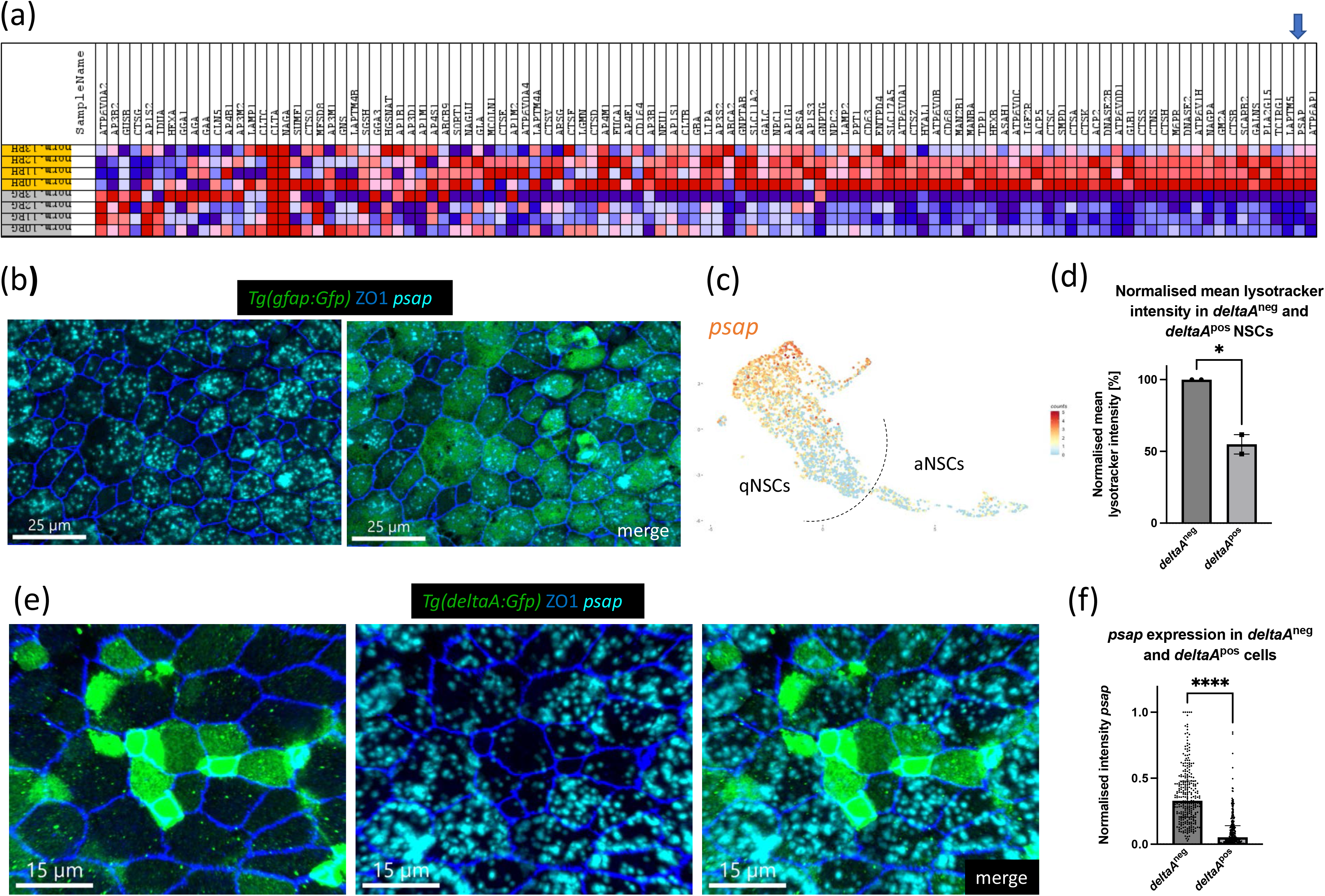
*psap* is heterogeneously expressed in quiescent NSCs. (**a**) Gene expression heatmap of the lysosome KEGG geneset (red = upregulated, blue = downregulated). Each column represents one biological replicate (grey column = aNSCs, yellow column = deeply qNSCs – *her4*^high^) – based on our bulk RNASeq dataset. Arrow points to *psap*. **(b)** Whole-mount RNAScope for *psap* (cyan) and immunostaining for *Tg(gfap:eGFP)* (NSCs, green) and ZO-1 (tight junctions, blue). Dorsal view of the pallial ventricular zone at 3 mpf. *psap* is heterogeneously expressed. Scale bar: 25 µm. **(c)** Umap of *psap* expression in NSCs based on the scRNA-seq dataset of [38]. *psap* is enriched in cells with deepest/longest quiescence (compare with Fig.S4a). **(d)** Normalized mean lysotracker intensity in *deltaA*^pos^ and *deltaA*^neg^ NSCs; barplot with SD. Each dot represents one independent experiment with 10 pooled zebrafish brains. Unpaired t-test, p = 0.0218. **(e)** Whole-mount RNAScope for *psap* (cyan) and immunostaining for GFP (transcriptional reporter for *deltaA*, green) and ZO-1 (tight junctions, blue). Dorsal view of the pallial ventricular zone at 3 mpf. *psap* is expressed more strongly in *deltaA*^neg^ NSCs. Scale bar: 15 µm. **(f)** Mean normalized intensity sum of *psap* after 2D projection in *deltaA*^neg^ and *deltaA*^pos^ cells. The highest intensity value has been set to 1. Every dot represents one cell. 3 independent experiments. *psap* is expressed at higher levels in *deltaA^neg^* NSCs. Mann-Whitney test, p = <0.0001.

To assess the relationship between *psap*, the number of lysosomes and qNSC states associated with different quiescence depth/length, we first tested whether *deltaA* expression already correlates with distinct metabolic states signed by NSCs harboring more or fewer lysosomes. We used double transgenic Tg(*gfap:dtomato*);Tg(*deltaA:egfp*) fish, allowing us to dissociate dTomato^pos^,GFP^pos^ *deltaA*^pos^ NSCs and dTomato^pos^ GFP^neg^ *deltaA*^neg^ NSCs for flow cytometry analysis after lysotracker staining (Figure S4d). When analyzing the mean intensity of lysotracker fluorescence, we confirmed that *deltaA*^pos^ NSCs contain fewer lysosomes than *deltaA*^neg^ NSCs (Median and IQR: 54.93 %, 48.17-61.69 % in *deltaA*^pos^ NSCs) (Figure 2d). We therefore analyzed *psap* expression in regard to *deltaA* by combining immunostainings for ZO1 (tight junctions) and GFP in transgenic Tg*(deltaA:egfp)* fish with RNAScope for *psap* (Figure 2e). We found that *psap* expression was significantly higher in *deltaA*^pos^ (median and IQR: 0.33, 0.21-0.48) compared to *deltaA*^neg^ (median and IQR: 0.05, 0.02-0.14) progenitors (Figure 2f). These results point to increased *psap* expression in qNSCs with deep/long quiescence, as opposed to qNSCs with shallower/shorter quiescence.

### 3.3. psap maintains NSCs in a state of deep quiescence

To explore the function of *psap* in quiescence *in vivo*, we chose a loss-of-function approach targeting individual NSCs in a salt-and-pepper fashion in the adult zebrafish pallium, allowing us to focus on cell-autonomous effects. Due to the duration of quiescence in adult pallial NSCs (doubling times on average of 124 days for *deltaA^neg^* and 28 days for *deltaA^pos^* NSCs) [11], we needed a method that could conditionally abrogate Psap function over several days/weeks while allowing cell tracking. Thus, we adapted and validated for the adult brain a recent RNAi knock-down method allowing long-term chase and concomitant tracing [48]. We designed 9 shRNAs targeting the endogenous *psap* transcript and 3 control shRNAs and cloned them in 3’ of pCMV:*h2amCherry*. Following electroporation into adult NSCs *in vivo*, we first verified that pCMV:*h2amCherry-RNAi(psap)* NSCs had a strong reduction of *psap* transcript levels compared to surrounding non-electroporated cells at 3 and 14 dpe, as measured using RNAscope (median and IQR: for 3d control 114.2 %, 113.0-115.4 % vs. RNAi(*psap*) 24.92 %, 13.73-28.62 %; for 14 d control 137.9 %, 130.4-145.3 % vs. RNAi(*psap*) 32.59 %, 21.07-40.00 %) (Figures S5a-d). We then proceeded to use this construct for loss-of-function experiments and electroporated Tg(*gfap:egfp*) fish to analyze H2amCherry^pos^,GFP^pos^ NSCs at 3, 5 and 14 days after knockdown (Figures 3a, b).

**Figure 3.**
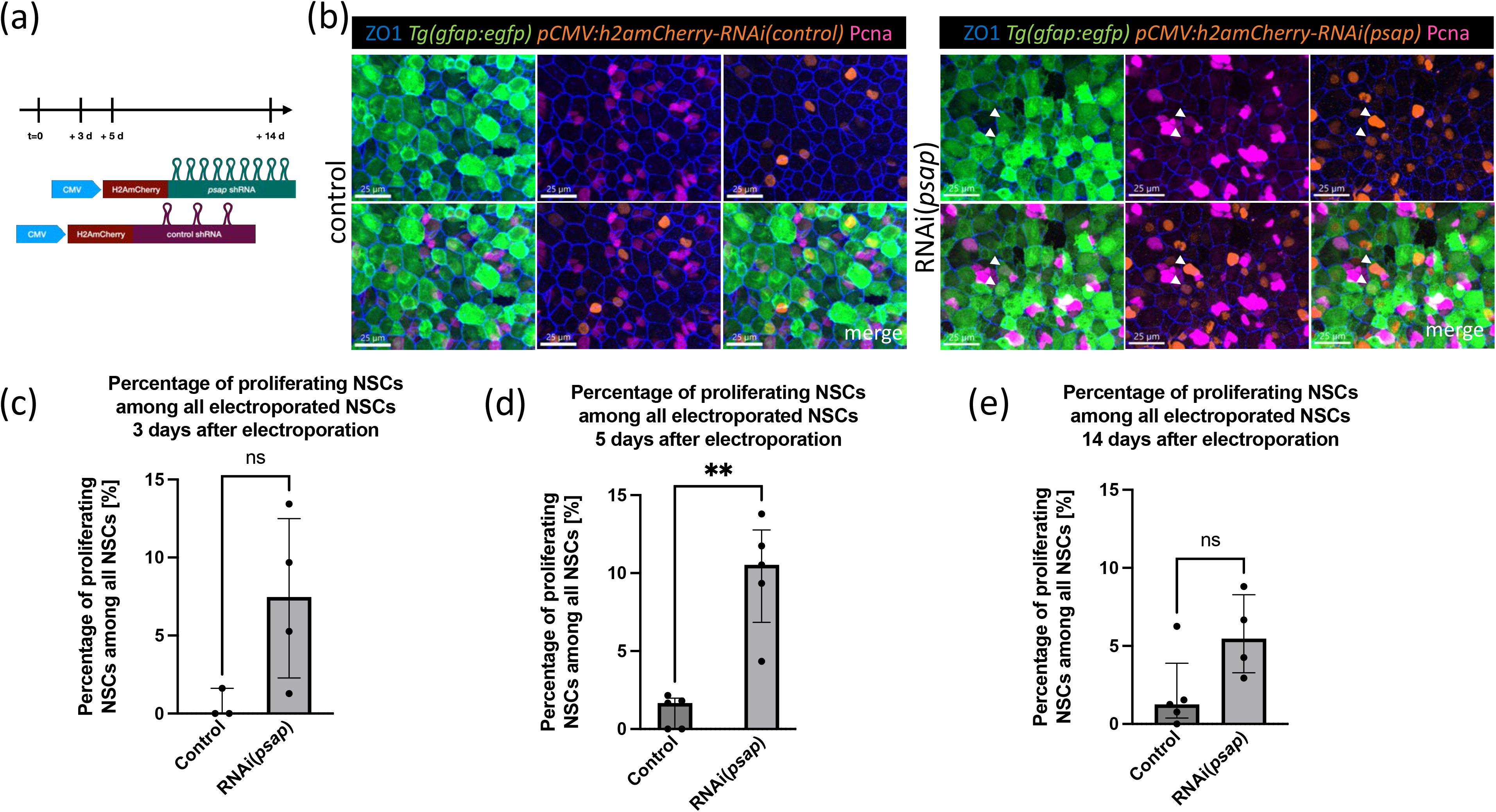
*psap* activity promotes NSC quiescence. (**a**) Experimental design: at day 0, *pCMV:RNAi(psap)* and *pCMV:RNAi(control)* were electroporated in pallial NSCs in 3 mpf fish. The fish were sacrificed at 3, 5 and 14 dpe and brains were processed for immunostaining. **(b)** Whole mount immunostaining for ZO1 (tight junctions, blue), GFP (NSCs, green), H2amCherry (electroporated cells, orange), and Pcna (proliferation, magenta) at 5 dpe in fish electroporated with the RNAi control construct vs RNAi(*psap*). Dorsal view of the pallium. Scale bar: 25 µm. Arrows point to proliferating NSCs, which are more frequent after loss of *psap*. Scale bar: 25 µm. **(c)** Percentage of aNSCs among all electroporated NSCs at 3 dpe in control– vs *psap*-RNAi electroporations. Each dot represents one animal. Line at median with IQR. Mann-Whitney test, p = 0.1143. **(d)** Percentage of aNSCs among all electroporated NSCs at 5 dpe in control– vs *psap*-RNAi electroporations. Each dot represents one animal. Line at median with IQR. Mann-Whitney test, p = 0.0079. *psap* RNAi increases the percentage of aNSCs significantly. **(e)** Percentage of activated NSCs among all electroporated NSCs at 14 dpe in control– vs *psap*-RNAi electroporations. Each dot represents one animal. Line at median with IQR. Mann-Whitney test, p = 0.0635.

Activation events were quantified by equaling each Pcna^pos^ doublet (sister cells post-division), and each Pcna^pos^ singlet, to one event. We observed a significantly increased proportion of H2amCherry^pos^,GFP^pos^,Pcna^pos^ aNSC events at 5 dpe upon *psap* knock-down. At 3 and 14 dpe we observed an increase, but this did not reach significance (median and IQR: 3d, control 0.00 %, 0.00-1.61 % vs. RNAi(*psap*) 7.47 %, 2.28-12.49 %; 5 d control 1.67 %, 0.00-1.98 % vs. RNAi(*psap*) 10.53 %, 6.85-12.77 %; 14 d control 1.25 %, 0.38-3.89 % vs. RNAi(*psap*) 5.46 %, 3.27-8.27 %) (Figures 3c-e). These results indicate that *psap* knockdown increases NSC entry into the cell cycle. Monitoring the ratio of Pcna^pos^ doublets over singlets further indicated that the cell cycle per se was not visibly affected (median and IQR: 3d, control 0.0 %, 0.0-0.0 %, vs. RNAi(*psap*) 11.61 %, 0.25-30.56 %; 5 d control 0.00 %, 0.00-33.33 % vs. RNAi(*psap*) 20.00 %, 5.57-33.33 %) (Figure S5e,f). Hence, loss-of-function experiments of *psap* increase NSC proliferation *in vivo*, identifying Psap as a novel factor necessary for NSC quiescence.

### 3.4. Increased NSC activation upon psap abrogation is, in part, associated with a progression along the NSC lineage hierarchy

Quiescence depth/duration, the successive steps from quiescence to activation, and NSC lineage progression, are intertwined processes on which Psap could act, with a common output measured by Pcna expression. For example, the duration of the *ascl1a^pos^* pre-activation state could be modified. Additionally, NSC quiescence and fate dynamics vary along lineage progression: self-renewing *deltaA^neg^* NSCs in deep/long quiescence generate, upon division, *deltaA^pos^* NSCs of shallow/short quiescence, characterized by stochastic fates biased towards the generation of neurons [11,37]. Thus, the increased NSC proliferation observed upon Psap abrogation can be interpreted as a general release of quiescence, a shortened pre-activation phase, a progression along the lineage, or combinations of these events.

To address this issue, we monitored the expression of *ascl1a* and *deltaA* upon loss of Psap function. Using RNAscope to quantify *ascl1a* expression at the single cell level *in situ,* in Tg(*gfap:egfp*) fish to identify NSCs (Figure 4a), we observed a significant increase in the proportion of GFP^pos^, *ascl1a*^pos^ NSCs 5 days post *psap* knockdown (median and IQR: control 9.41 %, 8.00-10.82 % vs. RNAi(*psap*) 25.00 %, 19.27-36.94 %) (Figure 4b), indicating a shift towards the pre-activated and/or committed state.

Next, we knocked down *psap* in Tg(*deltaA:egfp*) fish and monitored *deltaA* expression in electroporated cells. We observed an increase in the proportion of H2amCherry^pos^, GFP^pos^ cells at 5 dpe upon RNAi, from 22.97 %, 21.43-27.88 % (control RNAi, median and IQR) to 42.35 %, 37.82-51.71 % (*psap* RNAi, median and IQR) (Figures 4c and 4d). Despite the increased number of proliferating cells induced by *psap* RNAi (median and IQR: control 5.26 %, 2.58-6.58 vs. RNAi(*psap*) 9.73 %, 6.75-14.35 %) (Figure S6a), the proportion of proliferating cells among *deltaA*^pos^ progenitors stayed constant (median and IQR: control 16.67 %, 6.92-25.00 % vs. RNAi(*psap*) 21.86 %, 11.07-25.89 %) (Figure S6b). A similar conclusion was reached regarding the proportion of proliferating cells among *deltaA*^neg^ progenitors –although the low proliferation rate of *deltaA*^neg^ NSCs makes this analysis difficult-(median and IQR: control 0.00 %, 0.00-0.00 % vs. RNAi(*psap*) 0.00 %, 0.00-1.28) (Figure S6c). Together, these observations suggest that loss of *psap* leads to an initial shift towards the *deltaA^pos^* state, within which the transition towards activation and/or the proliferative behavior is not changed *per se*.

### 3.5. Neuronal generation is preserved upon psap abrogation

Finally, we aimed to address whether the fate of NSCs is altered after loss of *psap*. We first analyzed the fate of sister cells upon *psap* knock-down relative to *deltaA* expression: *deltaA^neg^* NSCs systematically self-renew upon division, while *deltaA^pos^* NSCs progressively engage towards neurogenesis, identifying different stemness potential [49]. Control and RNAi(*psap*) were electroporated into Tg(*deltaA:egfp*) fish and the relative distribution of division modes in doublets of sister cells was analyzed at 5 dpe. These distributions did not appear statistically different between control and RNAi(*psap*) electroporated cells (median and IQR: Mann-Whitney tests, *deltaA^neg^*/*deltaA^neg^* doublets: control: 0.00, 0.00-0.00, RNAi(*psap*): 6.25, 0.00-27.38, p = 0.3929; *deltaA^neg^*/*deltaA^pos^* doublets: control: 33.33, 0.00-100.00, RNAi(*psap*): 33.93, 0.00-70.00, p = 0.9524; *deltaA^pos^*/*deltaA^pos^* doublets: control: 66.67, 0.00-100.00, RNAi(*psap*): 41.43, 25.00-71.88, p = 0.6905).

To further substantiate this result, we analyzed the generation of more committed cell fates by focusing on terminal NSC divisions generating NPs (Gfap^neg^). For this, electroporations of RNAi(*psap*) into the Tg(*gfap:egfp*) line were used and NSC/NSC, NSC/NP and NP/NP doublets were counted. At 3, 5 and 14 dpe, we found that the relative distribution of NSC division modes was also similar between control and RNAi(*psap*) electroprated cells (at 5 dpe, median and IQR: Mann-Whitney tests, NSC/NSC doublets: control: 66.67, 8.33-29.17, RNAi(*psap*): 80.00, 41.67-81.53, p = 0.3929; NSC/NP doublets: control: 25.00, 0.00-100.00, RNAi(*psap*): 18.75, 11.69-30.83, p = 0.9524; NP/NP doublets: control: 16.67, 5.55-37.5, RNAi(*psap*): 9.091, 0.00-30.36, p = 0.6905) (Figures 4e) (see also Figure S7 for 3 and 14 dpe). Finally, we counted H2amCherry^pos^ cells at 14 dpe and calculated the ratio of neurons per NSC. We did observe large variations across samples in both control and *psap* RNAi samples and no significant difference between the control and knockdown conditions overall (median and IQR: control 0.2813, 0.1032-0.8405 vs. RNAi(*psap*) 0.4518, 0.3831-2.043) (Figure 4f).

**Figure 4:**
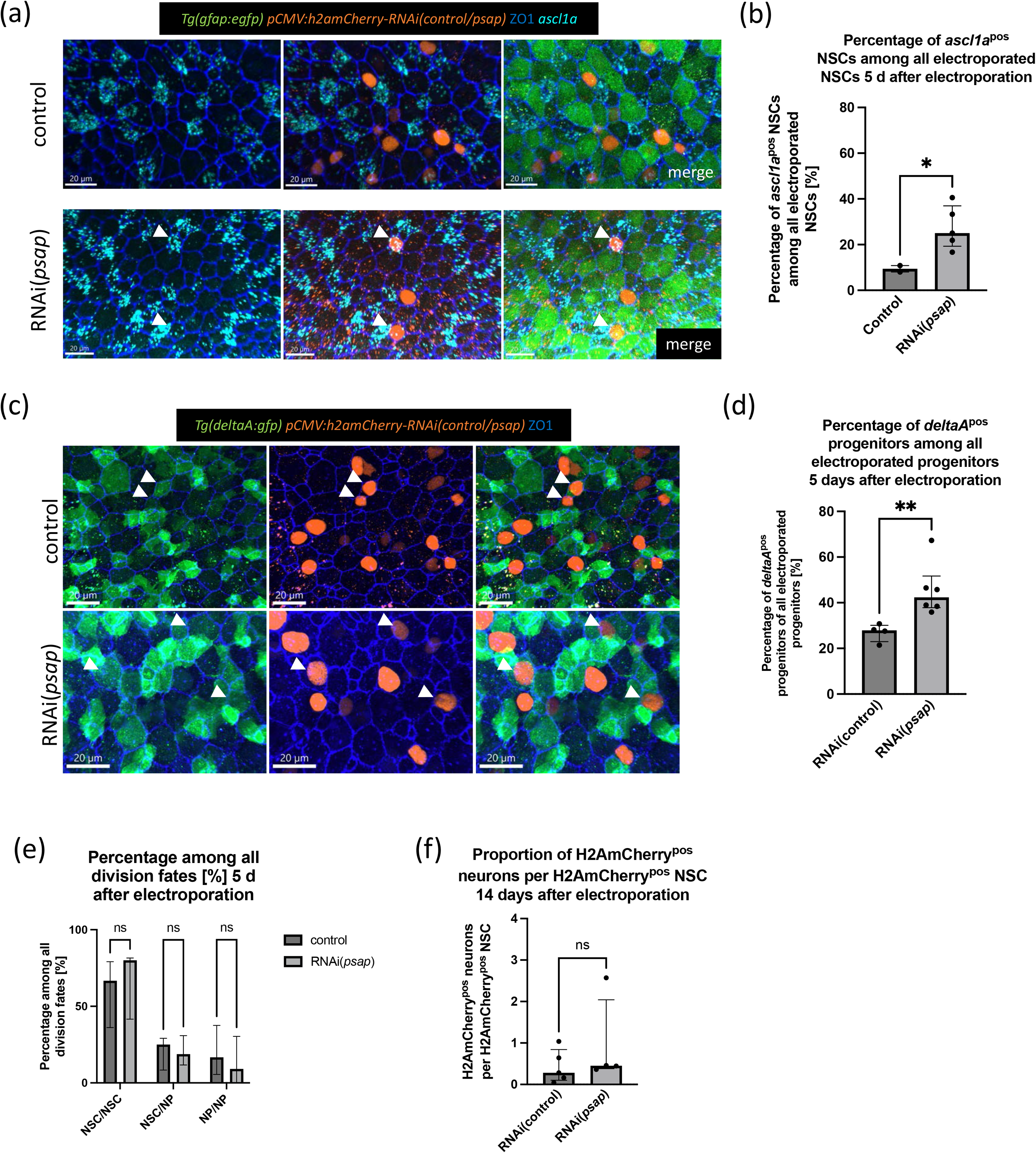
p*s*ap knock-down increases the proportion of NSCs in states associated with shallow quiescence, without affecting neurogenesis output. (**a**) RNAScope for *ascl1a* (turquoise) and whole mount immunostaining for gfap:GFP (green), H2amCherry (electroporated cells, orange) and ZO-1 (tight junctions, blue) 5 d after electroporation of control-(top) and *psap-*RNAi (bottom) construct in Tg(*gfap:egfp*) fish. Dorsal view of the pallial ventricular zone at 3 mpf. Arrows point to examples of *ascl1a*^pos^ cells. Loss of Psap leads to increased *ascl1a* expression. Scale bar: 20 µm. **(b)** Percentage of *ascl1a*^pos^ NSCs among all electroporated NSCs 5 d after control– and *psap*-RNAi electroporation. Line at median with IQR. Unpaired t-test with Welch’s correction, p = 0.0112. **(c)** Whole mount immunostaining for GFP (*deltaA*-expressing cells, green), H2amCherry (electroporated cells, orange), and ZO1 (tight junctions, blue) 5 days after electroporation of RNAi control (top) and RNAi(*psap*) construct. Dorsal view of the pallium at 3 mpf. Arrows point to GFP^pos^ progenitor cells. Scale bar: 20 µm. **(d)** Percentage of *deltaA*^pos^ progenitor cells among all electroporated progenitor cells 5 d after control– and *psap*-RNAi electroporation. Each dot represents one animal. Line at median with IQR. Mann-Whitney test, p = 0.0095. **(e)** Percentage of the different division modes in control vs RNAi(*psap)* electroporated cells at 5 dpe (see Fig.S7 for 3 and 14 dpe). x axis: state of the two sister cells in doublets. Lines at median with IQR. Mann-Whitney tests, NSC/NSC doublets: p = 0.8413; NSC/NP doublets: p = 0.7936; NP/NP doublets: p = 0.6429. **(f)** Ratio of neurons relative to NSCs in electroporated cells and their progeny. Each dot represents one animal. Line at median with IQR. Mann-Whitney test, p = 0.4127.

## 4. Discussion

This work brings novel molecular insight into the control of NSC quiescence in the adult brain. We show that lysosomal activity in adult zebrafish pallial NSCs correlates with and is needed for their quiescence. *In vivo*, we further show that, like lysosomal inhibition, blocking the sorting receptor Sortilin induces NSC proliferation. Finally, we provide evidence for the functional necessity of Prosaposin to maintain a NSC state associated with deep quiescence. These results together add support to the fact that lysosomes are positive regulators of NSC quiescence, likely through their role in autophagy, and show that proteins such as Prosaposin further refine this control to include the regulation of quiescent NSC substates associated with different quiescence depth or duration.

At the technical level, we made use of RNAi, a technique long debated in the zebrafish field due to nonspecific defects and high toxicity in embryos, likely due to a highjack of the microRNA machinery [50]. In 2015, efficient downregulation of mRNA through shRNAs was reported [48]. Adapting the system to single NSCs was successful in terms of knockdown efficiency, but also did not show any signs of toxicity, as assessed by NSC morphology and survival. The long-term perdurance of the induced knockdown with this system (at least up to 14 days), compared to morpholinos (3-4 days) [51], is advantageous, and it will be interesting to see whether other adult tissues are good targets for this method.

We show that qNSCs of the adult zebrafish pallium harbor more lysosomes and display a transcriptome enriched for genes associated with the endo-lysosomal compartment compared to their activated counterparts. Accordingly, the impairment of lysosomal function leads to an increase in NSC proliferation, which aligns with findings in the mouse SGZ [19] but not the SEZ [20]. Possible reasons for discrepancies between these mouse studies include their different neurogenic niches and experimental setups. The zebrafish adult pallial germinal niche appears, for its anterior component (Da-Dm, the focus of our *in situ* analyses), ontogenetically and molecularly related to the dorsal wall of the SEZ, itself closer to the SGZ in terms of markers expression (e.g., *Hopx*) [52], later entry into quiescence during development [53] and increased NSC quiescence depth [52,54], and cortical origin. The zebrafish adult pallial germinal niche also joins and includes the hippocampal area more postero-laterally. Like in [19], we found that concentrations of BafA at 50 nM led to toxicity. Lysosomal blockade can also drive cells into senescence [55], and it remains to be functionally tested whether any of these experimental setups increase senescence.

We further demonstrate that *psap* is highly expressed in deeply quiescent NSC. Psap is a prominent lysosomal protein (under its cleaved forms, Saposins, which hydrolyze sphingolipids) but can also be secreted, to boost lysosomal function in receiving cells [56] and/or act as a neurotrophic factor [57]. To center on the lysosomal function of Psap, we focused on cell-autonomous effects of *psap* manipulations. Specifically, the knockdown of *psap* shifted NSCs towards substates associated with shallower quiescence (as defined by *deltaA* and *ascl1a*), and with activation. The fact that lysosomal activity can modulate quiescence depth is an idea that was postulated in embryonic fibroblasts [58] and which is compatible with several studies in hematopoietic stem cells [59,60]. Although this was not directly addressed to date for NSCs, pre-existing studies highlighting progressive changes of NSC states are in line with our conclusion. For example, in cultured NSCs from the hippocampus, mRNA levels for TFEB or Cathepsins increase progressively during the first three days of incubation with the quiescence inducer BMP. Conversely, BafA leads to progressive increases in activated EGFR and Notch1-ICD, which is compatible with a transition from quiescence to activation through shallower quiescence stages [19].

Under physiological conditions, changes in NSC quiescence duration in the adult zebrafish pallium and mouse SGZ are concomitant with NSC progression towards a neurogenic fate [37,49,61]. Thus, a limitation to current studies, including ours, is to distinguish between a *bona fide* function of lysosomal factors (e.g., Psap) in quiescence depth/duration per se vs. in NSC position along the lineage (i.e. self-renewal vs an engagement towards a neurogenic fate). A primary effect on quiescence could secondarily affect neurogenesis commitment, and a primary effect on neurogenesis commitment could secondarily affect quiescence. In the future, it will be important to resolve whether *deltaA* and *ascl1a* expression induced under *psap* knock-down reflect decreased quiescence depth, increased commitment, or both. Verifying this point will require more extensive lineage tracing, as the number of RNAi(*psap*) NSCs in electroporation experiments is unfortunately too low for these cells to be profiled and their state characterized in depth.

A remaining question will be to identify the relevant targets of Psap. In the adult mouse SGZ, impaired lysosomal function leads to the accumulation of phosphorylated EGFR, Notch1-ICD and Cyclin D1 [19]. Our attempts to detect such changes in adult zebrafish pallial NSCs *in vivo* (using anti-EGFR IHC and a direct Notch3 signaling reporter) remained unsuccessful. We also were technically limited *in vivo* to address the hydrolysis of sphingolipids [43], or related changes in metabolism due to defective mitochondria and oxidative phosphorylation [62,63]. Mitochondrial function depends on intact lysosomes [59,60,64] and is important for hippocampal neurogenesis [65]. Thus, future directions could explore the link between lysosomes and mitochondria in NSCs.

## Supplementary Materials

Figure S1: *her4.3* expression and GSEA enrichment plots;

Figure S2: Gating strategy for lysotracker experiment;

Figure S3: Umaps of *sort1a* and *sort1b* expression in adult pallial NSCs;

Figure S4: Use of *deltaA* expression as a marker for qNSCs in with a state with shallow/shorter quiescence;

Figure S5: RNAi(*psap*) efficiently reduces *psap* RNA in transfected cells;

Figure S6: Characterization of the *deltaA^pos^* state upon *psap* knock down;

Figure S7: Percentage of the different division modes relative to *gfap*:GFP expression (NSCs: Gfap^pos^, NPs: Gfap^neg^) in control vs RNAi(*psap)* electroporated cells;

Table S1: Genes upregulated in RFP^high^ qNSCs vs aNSCs,

Table S2: Genes downregulated in RFP^high^ qNSCs vs aNSCs,

Table S3: Genes upregulated in 3 mpf vs 1.5 mpf qNSCs, Table S4: Genes downregulated in 3 mpf vs 1.5 mpf qNSCs.

## Author Contributions

Conceptualization, M.L. and L.BC.; methodology, M.L, M.T., E.TT, S.O., L.M.; validation, M.L.; investigation, M.L.; resources, D.M., M.C., H.V., R.L.; data curation, M.L., E.TT.; writing—original draft preparation, M.L.; writing—review and editing, all authors.; visualization, M.L.; supervision, L.BC.; project administration, L.BC.; funding acquisition, L.BC. All authors have read and agreed to the published version of the manuscript.

## Institutional Review Board Statement

The animal study protocol was approved by the Ethics Committee n°39 of Institut Pasteur (authorization #36936, April 26^th^, 2022) and DDPP-2021-021 of the Direction Départementale de la Protection des Populations de Paris.

## Informed Consent Statement

Not applicable.

## Data Availability Statement

the RNASeq data are available on GEO under accession number GSE239598 (https://www.ncbi.nlm.nih.gov/geo/query/acc.cgi?acc=GSE239598 using the following password: ghcfwwugtzavfsl).

## Supporting information

Graphical abstract

## Acknowledgments

We are greatly indebted to Jean Giacomotto and Alisha Tromp for their help with the RNAi protocol and for providing the 3’ entry backbone for these constructs. We thank Sébastien Bedu and Nicolas Dray for help in generating the Tg(*mcm5:nls-rfp*) line. We thank the ZEN team and Thomas Wollert for input, and the Institut Pasteur CB UTechS service platform for expert assistance with FACS analysis.

