## Supplementary figure legends for "Prosaposin maintains adult neural stem cells in a state associated with deep quiescence"

Labusch, M. et al.

### Supplementary Materials

Figure S1: *her4.3* expression and GSEA enrichment plots;

Figure S2: Gating strategy for lysotracker experiment;

Figure S3: Umaps of *sort1a* and *sort1b* expression in adult pallial NSCs;

Figure S4: Use of *deltaA* expression as a marker for qNSCs in with a state with shallow/shorter quiescence;

Figure S5: RNAi(*psap*) efficiently reduces *psap* RNA in transfected cells;

Figure S6: Characterization of the *deltaA*$^{pos}$ state upon *psap* knock down;

Figure S7: Percentage of the different division modes relative to *gfap*:GFP expression (NSCs: Gfap$^{pos}$, NPs: Gfap$^{neg}$) in control vs RNAi(*psap)* electroporated cells;

Table S1: Genes upregulated in RFP$^{high}$ qNSCs vs aNSCs,

Table S2: Genes downregulated in RFP$^{high}$ qNSCs vs aNSCs,

Table S3: Genes upregulated in 3 mpf vs 1.5 mpf qNSCs,

Table S4: Genes downregulated in 3 mpf vs 1.5 mpf qNSCs.

**Legends for supplementary figures**

**Figure S1: *her4.3* expression and GSEA enrichment plots. (a)** Umap of *her4.3* expression in adult pallial NSCs based on the scRNA Seq dataset of Morizet *et al.*, 2023 [xx]. The position of qNSCs vs aNSCs is indicated, compare with Fig. S4(a) for *pcna*. **(b-h)** GSEA enrichment plots from the KEGG pathway **(a)** "Cell Cycle", **(b)** "DNA Replication", **(c)** "Ribosome", **(d)** "Proteasome", **(e)** "Lysosome", **(f)** Regulation of Autophagy", **(g)** "Endocytosis". The enrichment score (ES) is represented on the y axis. On the x axis are genes from the given gene set, represented as black lines. The green line connects the ES with the genes. The ES is based on the strongest deviation from zero as calculated for each gene going down the ranked genelist, and represents the degree of overrepresentation of a gene set at the top or the bottom of the ranked gene list. The coloured band at the bottom represents the degree of correlation of genes with aNSCs (red) or qNSCs (blue). Cell cycle, DNA Replication, Ribosome and Proteasome genes were enriched in aNSCs, whereas genes associated with lysosomes, autophagy and endocytosis are enriched in qNSCs.

**Figure S2: Gating strategy for lysotracker experiment. (a)** Gating strategy for flow cytometry analysis of qNSCs (GFP$^{pos}$ RFP$^{neg}$) vs. aNSCs (GFP$^{pos}$ RFP$^{pos}$) from adult Tg(*gfap:egfp*);Tg(*mcm5:nls-rfp*) telencephali. **(b)** Umap of *sort1a* expression in NSCs based on the scRNA Seq dataset of Morizet *et al.*, 2023. **(c)** Umap of *sort1b* expression in NSCs based on the scRNA Seq dataset of Morizet *et al.*, 2023 [38].

**Figure S3:** Umaps of *sort1a* and *sort1b* expression in adult pallial NSCs based on the scRNA Seq dataset of Morizet *et al*., 2023 [38].

**Figure S4:** Use of *deltaA* expression as a marker for qNSCs in with a state with shallow/shorter quiescence. **(a)** Umaps of *pcna, ascl1a* and *deltaA* expression in NSCs based on the scRNA Seq dataset of Morizet *et al*., 2023 [38]. **(d)** Gating strategy for flow cytometry analysis of *deltaA*$^{neg}$ NSCs (GFP$^{neg}$ dTomato$^{pos}$) and *deltaA*$^{pos}$ NSCs (GFP$^{pos}$ dTomato$^{pos}$) from adult telencephala.

**Figure S5: RNAi(*psap*) efficiently reduces *psap* RNA in transfected cells. (a)** RNAScope for *psap* (cyan) and whole mount immunostaining for ZO1 (tight junctions, blue), H2amCherry (orange, electroporated cells) 3 d after electroporation of control- and *psap*-RNAi constructs. Dorsal view of the pallial ventricular zone at 3 mpf. Dotted white lines surround electroporated cells. Scale bar: 20 µm. **(b) N**ormalized mean fluorescence intensity of *psap* 3 d after control- and *psap*-RNAi electroporation. Each dot represents one animal. Line at median with IQR. Unpaired t-test with Welch's correction, p = <0.0001. RNAi against *psap* decreases *psap* RNA, thus validating the method. **(c)** RNAScope for *psap* (cyan) and whole mount immunostaining for ZO1 (tight junctions, blue), and H2amCherry (orange, electroporated cells) 14 d after electroporation of control- and *psap*-RNAi construct. Dorsal view of the pallial ventricular zone at 3 mpf. Dotted white line surrounds electroporated cells. Scale bar: 20 µm. **(d)** Mean normalized fluorescence intensity of *psap* 14 d after control- and *psap*-RNAi electroporation. Each dot represents one animal. Line at median with IQR. Unpaired t-test with Welch's correction, p = 0.0065. RNAi against *psap* decreases *psap* RNA, thus validating the method. **(e)** Percentage of divisions with cytokinesis among all electroporated NSCs 3 d after control- and *psap*-RNAi electroporation. Line at median with IQR. **(f)** Percentage of divisions with cytokinesis among all electroporated NSCs 5 d after control- and *psap*-RNAi electroporation. Line at median with IQR. Mann-Whitney test, p = 0.5.

**Figure S6: Proliferation of *deltaA*$^{pos}$ cells upon *psap* knock down. (a)** Percentage of proliferating progenitor cells among all electroporated progenitor cells 5 d after control- and *psap*-RNAi electroporation. Each dot represents one animal. Line at median with IQR. Mann-Whitney test, p = 0.0381. **(b)** Percentage of proliferating progenitor cells among all electroporated *deltaA*$^{pos}$ progenitor cells 5 d after control- and *psap*-RNAi electroporation. Each dot represents one animal. Line at median with IQR. Mann-Whitney test, p = 0.9286. **(c)** Percentage of proliferating progenitor cells among all electroporated *deltaA*$^{neg}$ progenitor cells 5 d after control- and *psap*-RNAi electroporation. Each dot represents one animal. Line at median with IQR. Mann-Whitney test, p > 0.9999.

**Figure S7:** Percentage of the different division modes relative to *gfap*:GFP expression (NSCs: Gfap$^{pos}$, NPs: Gfap$^{neg}$) in control vs RNAi(*psap)* electroporated cells. x axis: state of the two sister cells in doublets. Lines at median with IQR. Mann-Whitney tests. **(a)** at 3 dpe. NSC/NSC doublets: control: 33.33, 0.00-75.00, RNAi(*psap*): 73.33, 54.17-8429, p = 0.2286; NSC/NP doublets: control: 44.44, 12.50-100.00, RNAi(*psap*) 13.39, 10.63-2857, p = 0.2857; NP/NP doublets: control: 12.50, 0.00-22.22, RNAi(*psap*): 5.00, 0.00-30.63, p = 0.9143. **(b)** at 14 dpe. NSC/NSC doublets: control: 77.78, 56.06-85.56, RNAi(*psap*): 54.55, 18.86-75.91, p = 0.3413; NSC/NP doublets: control: 9.09, 3.33-13.89, RNAi(*psap*): 16.36, 2.50-32.95, p = 0.4524; NP/NP doublets: control: 11.11, 2.78-39.39, RNAi(*psap*): 20.91, 2.5-75.45, p = 0.9762.
