## supplementary figures for "Prosaposin maintains adult neural stem cells in a state associated with deep quiescence"

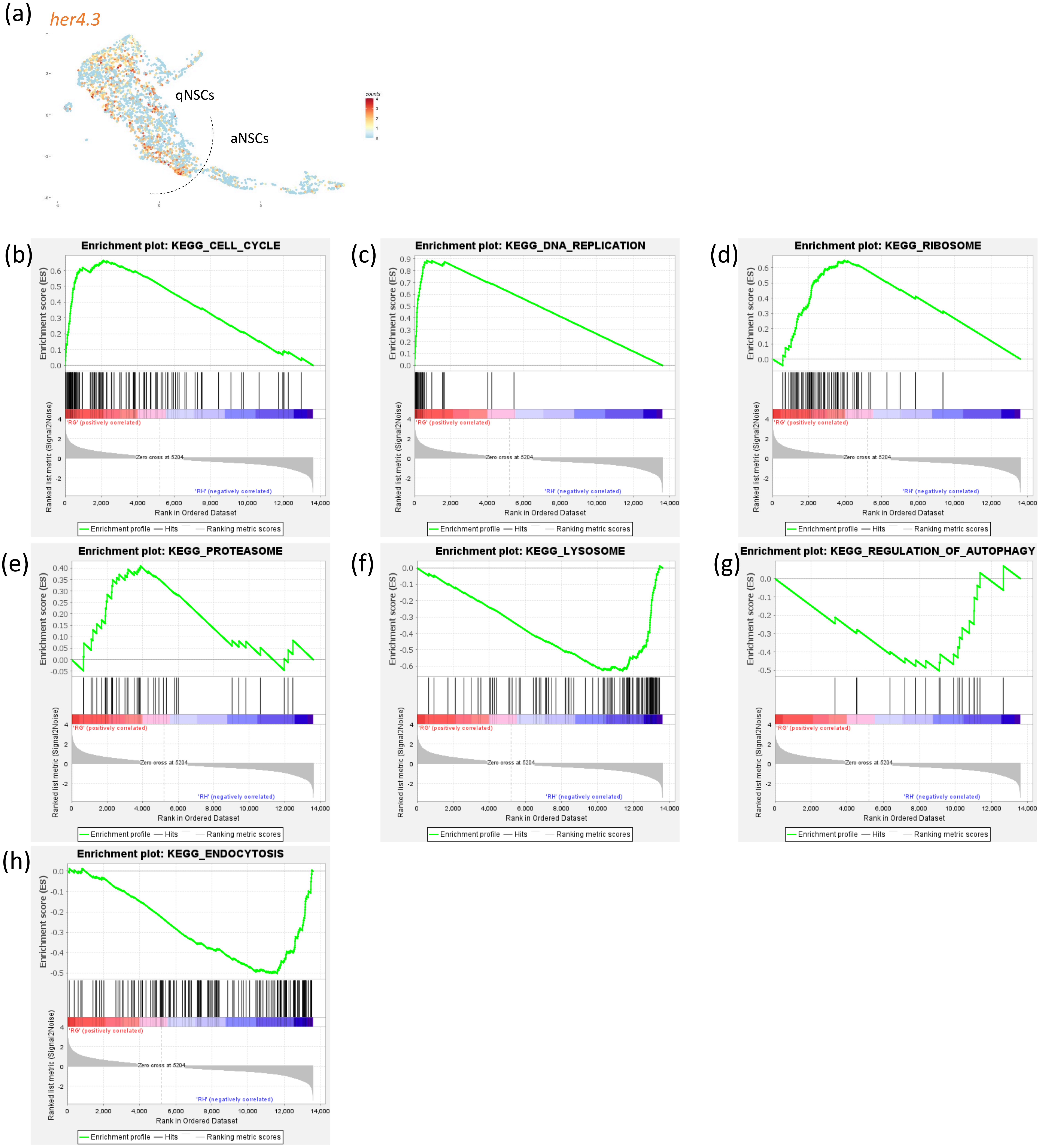


Labusch et al., Figure S1

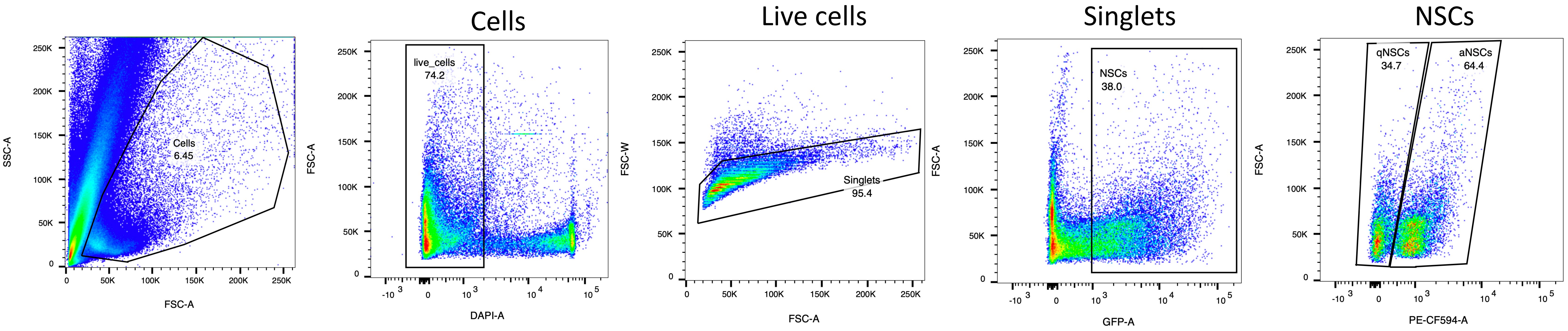


Labusch et al., Figure S2

(b) *sort1a* (c) *sort1b*

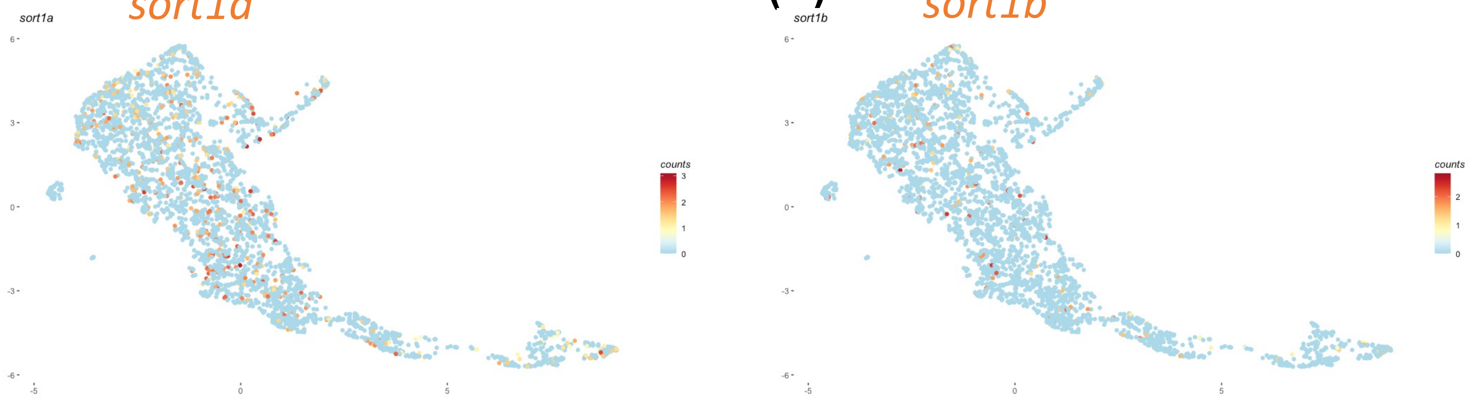


Labusch et al., Figure S3

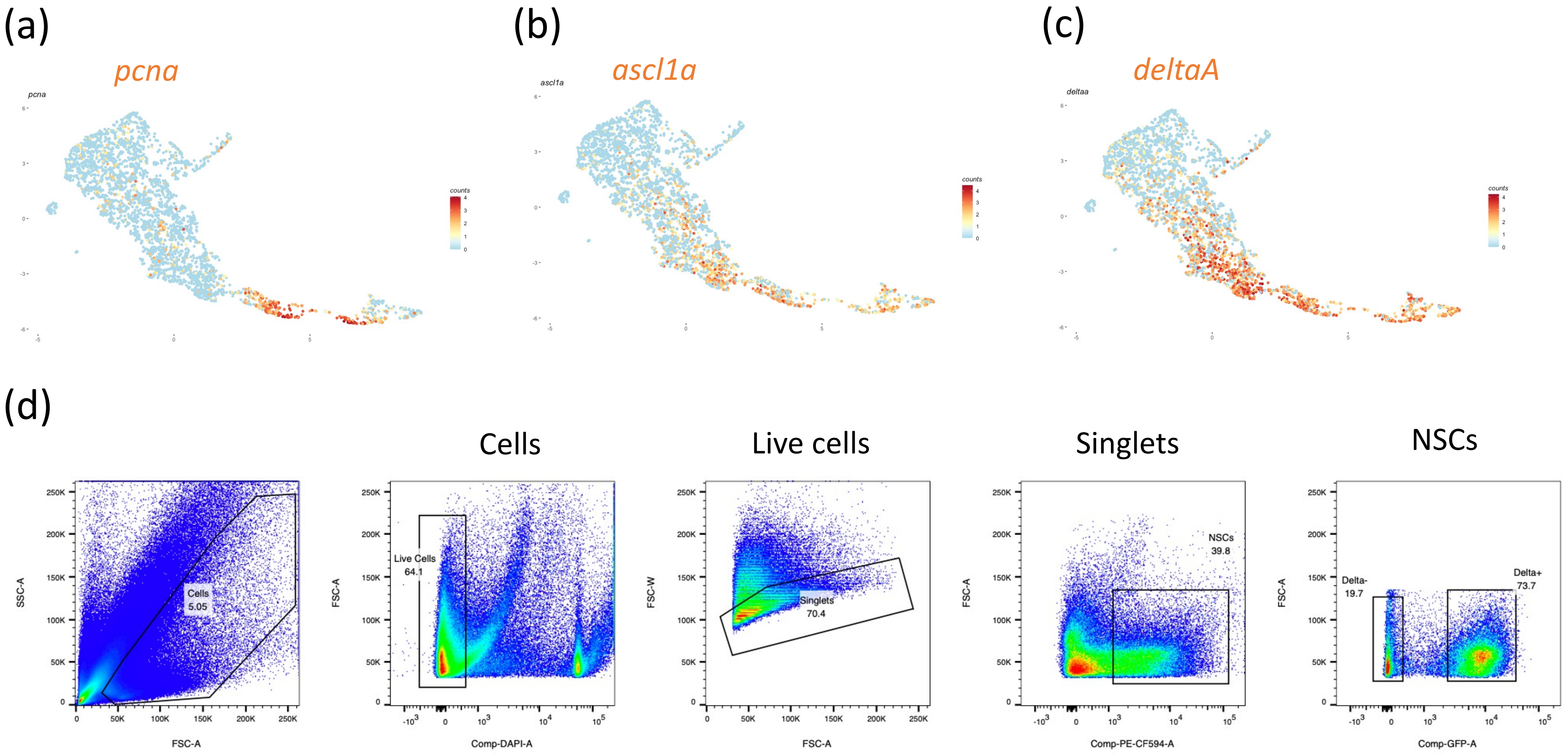


Labusch et al., Figure S4

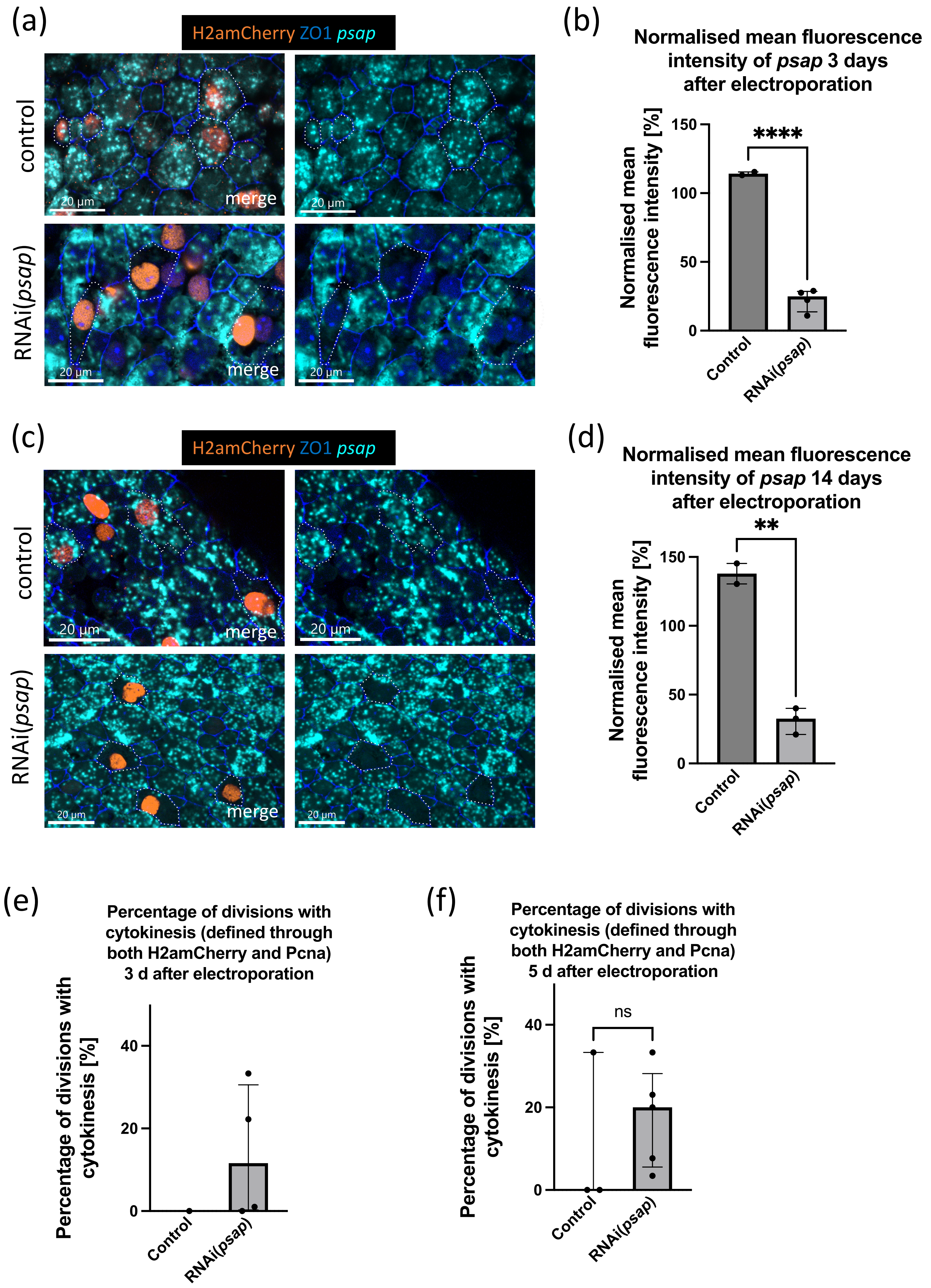


Labusch et al., Figure S5

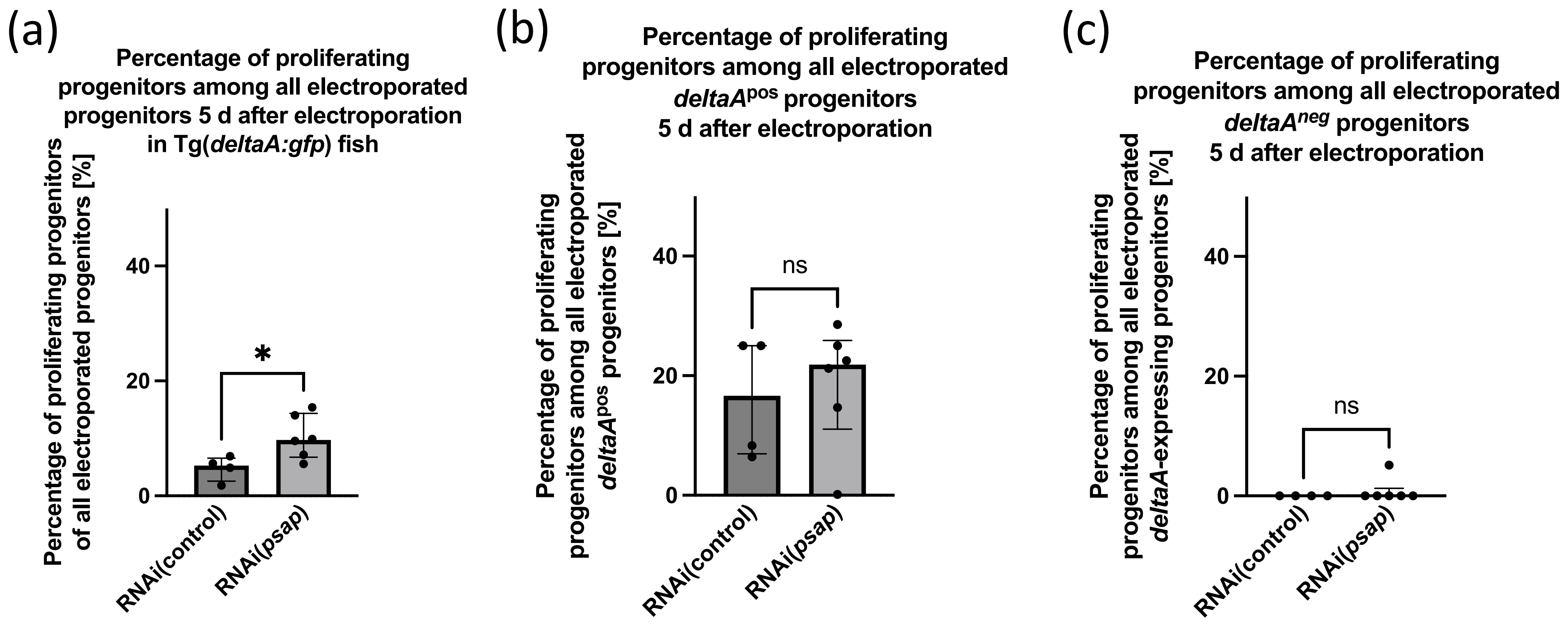


Labusch et al., Figure S6

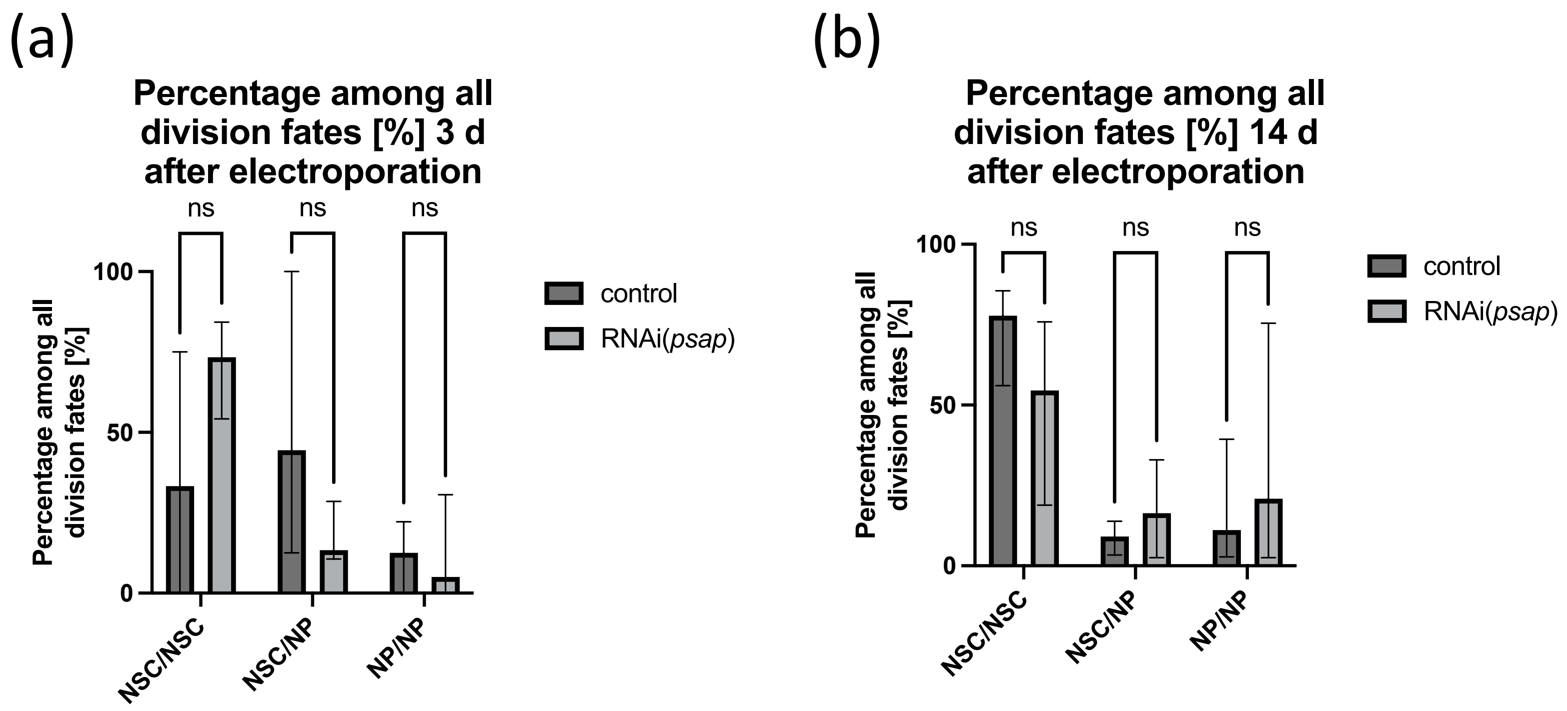


Labusch et al., Figure S7
