## Supplementary figures and images for "Prosaposin maintains adult neural stem cells in a state associated with deep quiescence"

### Graphical abstract

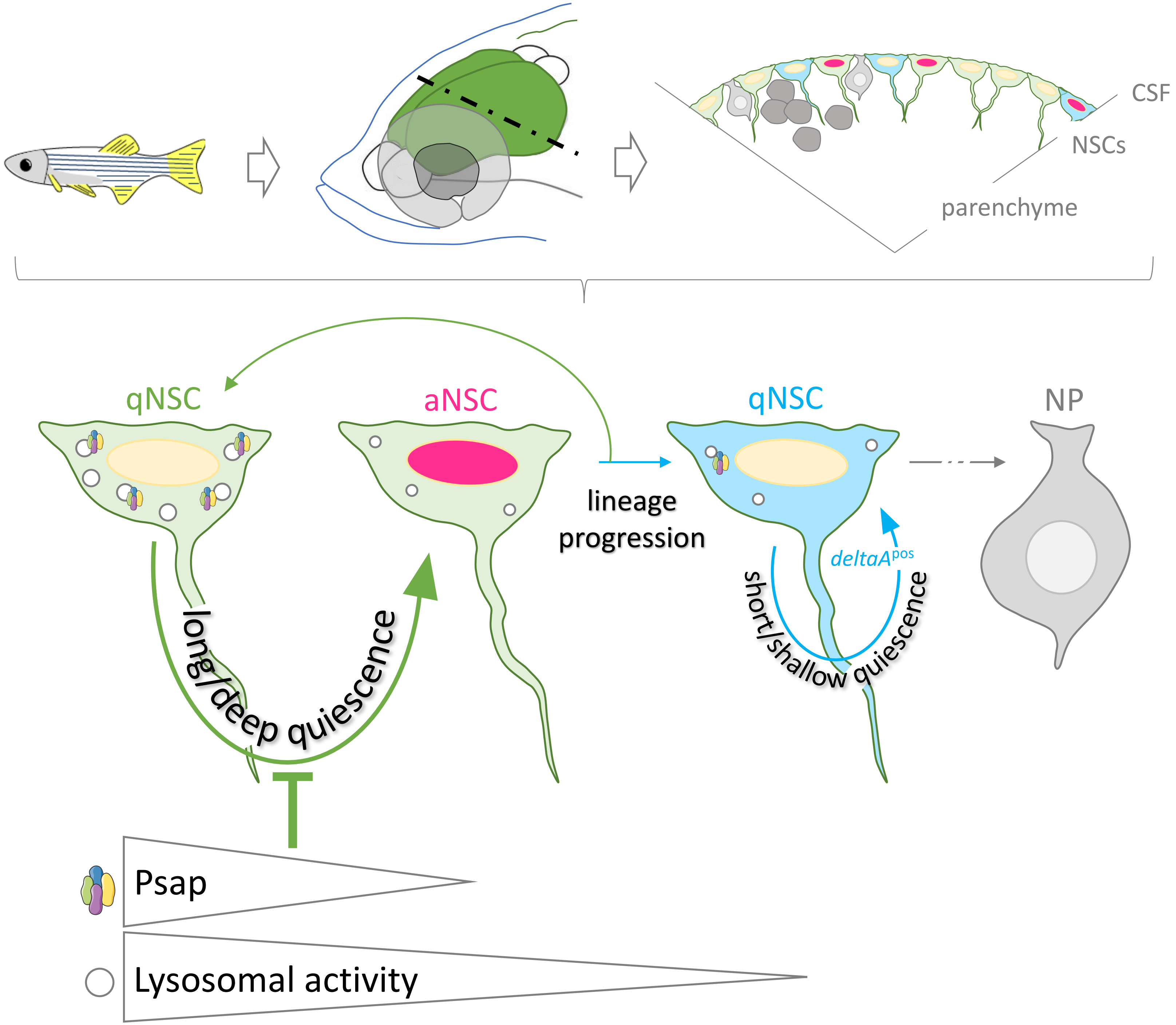
